# Loss of immune signalling pathways and increased pathogen susceptibility associated to photosymbiosis in acoels

**DOI:** 10.1101/2025.06.17.660074

**Authors:** Francesca Pinton, Nadezhda N. Rimskaya-Korsakova, Katja Felbel, Elisabeth Grimmer, Andreas Hejnol

## Abstract

**Background:** Host immunity plays an important role in coral symbiosis with dinoflagellates. Photosymbiosis (the association between animal hosts and photosynthetic endosymbionts) has evolved multiple times within animals, *e.g.* within acoels, which are soft-bodied marine invertebrates whose immunity remains so far undescribed.

**Results:** Our predicted proteome searches show that acoels lack immune signal transduction genes commonly conserved in animals. Their loss in acoels predates the occurrence of photosymbiosis in this clade. Immune challenges with the coral pathogen and bleaching agent, *Vibrio coralliilyticus,* increase acoel mortality and decrease symbiont numbers in adults of the photosymbiotic acoel *Convolutriloba macropyga.* Mortality in aposymbiotic *C. macropyga* juveniles or aposymbiotic species *Hofstenia miamia* is not affected. Ultrastructural studies of immune challenged animals by Transmission Electron Microscopy show damages at the cellular and organelle level, as well as a degradation of potential pathogens by the host. *In situ* hybridisation and differential gene expression analysis point to some areas of interaction between Pattern Recognition Receptors and microbes, as well as to the involvement of acoel-specific or uncharacterised genes.

**Conclusions:** Based on our findings, photosymbiosis evolution in acoels could have been favoured by the loss of immune signalling pathways. Photosymbiosis in acoels seems to increase susceptibility to pathogen exposure and is disrupted by pathogens. Our data also suggests the possibility of a novel molecular immune response specific to acoels.

## Background

Photosymbiosis is the association between an animal host and endosymbionts capable of photosynthesis [1]. It is usually considered a mutualism, with the animal host providing protection and inorganic compounds, while receiving photosynthates from the endosymbionts [1–4]. The best studied example of photosymbiosis is the one between cnidarians—corals or sea anemones—and dinoflagellates. This association is crucial for the coral reef ecosystem and increasingly endangered by climate change [5–10]. Increasing water temperatures, ocean acidification, and other stressors bring to a disruption of the coral-dinoflagellate photosymbiosis (*i.e.* dysbiosis) [11–17], with loss of dinoflagellate pigments or loss of the dinoflagellates themselves. Symbiont loss can happen by degradation, expulsion by the host, detachment or death of the animal cell hosting them [18–20]. Dysbiosis in corals is associated with the appearance of white spots (bleaching), increased mortality, and susceptibility to diseases [21–23]. Bacterial pathogens can also cause coral bleaching, as well as tissue lysis [21,24–26]. Research on the immune system of cnidarians has yielded important insights on the establishment, maintenance, and disruption of cnidarian photosymbiosis [1,19,27], as well as on the evolution of innate immunity [28–31].

Cnidarians are not the only animals hosting photosynthetic endosymbionts. Photosymbiosis is a widespread phenomenon in animals and has evolved several times, involving a great variety of photosynthetic partners [1,3,4]. Addressing photosymbiosis and its link to immunity in an evolutionary context is crucial to understand its underlying mechanisms and to make accurate predictions. To this end, we need to expand the pool of photosymbiotic systems studied [1,32].

Acoela (Xenacoelomorpha) are flat soft-bodied bilaterians, mostly found in marine habitats [33–35]. Some acoel species rely exclusively on photosymbiotic endosymbionts for nutrition (*e.g. Symsagittifera roscoffensis*), others regularly feed but are still dependent on their symbionts (*e.g. Convolutriloba macropyga*), and some others do not establish photosymbiotic relationships at all (*e.g. Hofstenia miamia*) [36,37]. Multiple photosymbiotic acoel species inhabit tropical reefs, alongside corals [38]; some are even considered parasitic to corals [39–42]. An increase in temperature causes symbiosis disruption and mortality in *Convolutriloba* species [37,43]. Water acidification, on the contrary, does not cause mortality in the acoel *S. roscoffensis*, and it only leads to bleaching by symbiont expulsion if extremely high [44]. Acoel responses to pathogens, including bleaching-inducing pathogens, remain uncharacterized. Furthermore, photosymbiosis interaction with the immune system has yet to be understood in this clade.

Here, we investigate the relationship between photosymbiosis and the immune system in acoels. First, our findings on xenacoelomorph immune gene repertoire and photosymbiosis presence are presented in an evolutionary context. Then, we use *in vivo* immune challenges to characterize acoel responses to the cnidarian pathogen and bleaching agent *Vibrio coralliilyticus.* We find an increased mortality in photosymbiotic acoels, but not in non-photosymbiotic ones. In photosymbiotic acoels, tissue damage can be observed, as well as symbiosis disruption. Our investigation of molecular responses to pathogens suggests the possibility of completely novel immune mechanisms in acoels.

## Results

### Photosymbiosis likely evolved twice within Acoela

To investigate the coevolution of photosymbiosis and the immune system in acoels, we started by mapping the presence and type of photosynthetic endosymbionts on the phylogenetic tree of acoels and their closest relatives (Fig.1A, Table S1). We followed the most robust Xenacoelomorpha phylogeny to date [45]. Since it does not feature many species of Convolutidae—the acoel clade containing all photosymbiotic species—we followed Jondelius *et al.* [46] for phylogenetic relationships within Convolutidae. Photosynthetic endosymbionts can be green algae (Chlorophyta), dinoflagellates (Dinoflagellata), or diatoms (Bacillariophyceae) [36]. All photosymbiotic acoels belong to Convolutidae and cluster compactly in two groups: (1) species with green algae as endosymbionts and *Convoluta convoluta*–the only species with diatom symbionts–, (2) species in symbiosis with dinoflagellates. A third Convolutidae clade, sister to (1), only contains non-photosymbiotic species.

**Fig. 1.**
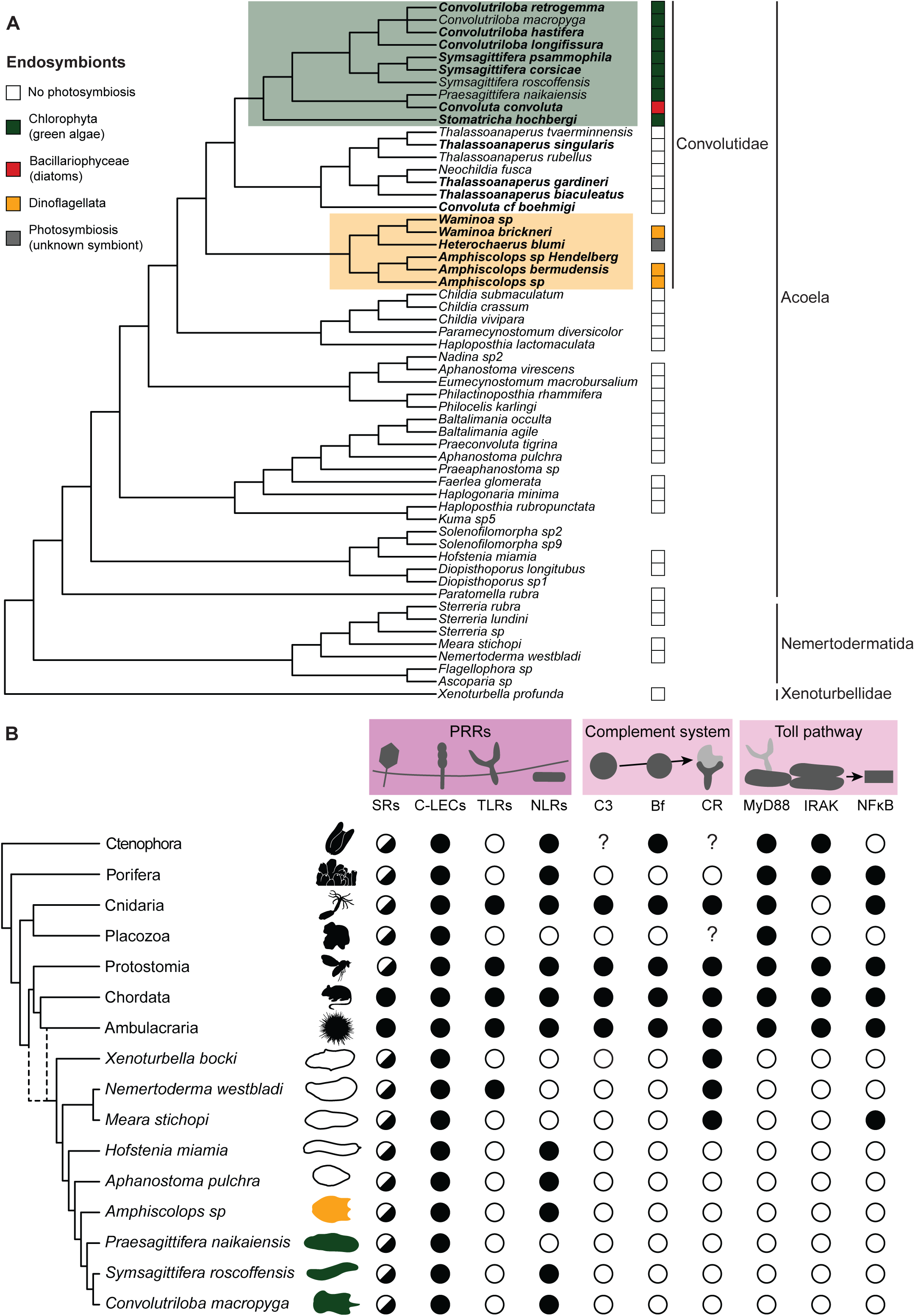
Photosymbiosis and immune genes in Xenacoelomorpha. **A** Photosynthetic endosymbiont presence and type mapped on the phylogenetic tree of Xenacoelomorpha. Tree after Abalde & Jondelius [45], relationships between species in bold after Jondelius *et al.* [46]. No square means no data available. Presence and type of photosynthetic endosymbionts for each species can be found in Table S1, together with references for each observation. Some silhouettes are from Phylopic (www.phylopic.org – credits to Andreas Hejnol, Soledad Miranda-Rottermann, Noah Schlottman, Michelle Site, Marina Vingiani, Jake Warner). **B** Immune genes in Xenacoelomorpha and other animal groups: black = present; white = absent; half circle = only some types present; question mark = no data. Metazoan tree after Dunn *et al.*, Laumer *et al.*, Schultz *et al.* [47–49]. Sources for gene presence/absence in non-xenacoelomorph metazoans: Kamm *et al.* [50] for all genes in Placozoa; Orús-Alcalde *et al.* [51] for TLR; Song *et al.* [52] for Toll pathway in Porifera, Cnidaria, Protostomia, Chordata; Orús-Alcalde *et al.* [53] for Toll pathway and complement system in Porifera, Cnidaria, Protostomia, Chordata, Ambulacraria; Rathinam *et al.* [31] for NLRs in Porifera, factor B in Ctenophora, SRs, C-lectins, NLR in Ambulacraria; Zelensky and Gready [54] for C-lectins in Porifera, Cnidaria, Protostomia, Chordata; Neubauer *et al.* [55] for SRs in Cnidaria and Chordata; Pancer *et al.* [56] for SRs in Porifera; Canton *et al.*, Melo Clavijo *et al.* [57,58] for SRs in Protostomia; Zhu *et al.* [59] for NLRs in Ctenophora, Porifera, Cnidaria, Protstomia, Ambulacraria, Chordata; Koutsouveli *et al.* [60] for PRRs, MyD88 in Ctenophora; Traylor-Knowles *et al.* [61] for IRAK and NFkB in Ctenophora. For Ctenophora, we preferred gene presence/absence inferred from domain searches and using genomic data [60,61].

These three clades are well-supported in the original studies, with Ultrafast bootstrap/SH-like approximate likelihood ratio test > 95 [45] or posterior probability > 0.90 [46]. This supports the claim that photosymbiosis evolved twice in acoels [38], always within Convolutidae, but once with dinoflagellates and once with green algae or diatoms (clades highlighted in Fig. 1A).

### Pattern Recognition Receptors are mostly conserved in acoels, while innate immune signalling pathways are absent

To characterize the acoel immune system, we first searched for innate immune genes in available Xenacoelomorpha genomes and transcriptomes (Fig. 1B). We focused on genes that are conserved in metazoans and involved in pattern recognition and signal transduction, especially the ones important for cnidaria-dinoflagellate symbiosis [1,19,27,62]. Since immune genes are often under rapid directional selection [63,64] and acoels are fast evolving [45], we searched for predicted proteins based on the presence of conserved domains in the expected order (Fig. S1), and, when relevant, based on their phylogenetic relationships (Fig. S2-5).

Pattern Recognition Receptors (PRRs) are responsible for the first interaction with microbes [65]. We searched for domain patterns corresponding to Scavenger receptors (SRs), C-type lectins, Toll-like receptors (TLRs), and NOD-like receptors (NLR) (Fig. S1A). SRs are characterized by a class-specific domain structure and not by phylogenetic relatedness [55,57,58,66–68]. No predicted protein in the xenacoelomorph species investigated shows domain combinations characteristic of SR class A. SR class B are found in all investigated species, although *Nemertoderma westbladi* sequences have only one transmembrane region instead of 2 (Fig. S1A). SR class E are found in all investigated species and SR class I only in the xenoturbellid *Xenoturbella bocki* and in the acoels *Hofstenia miamia* and *Aphanostoma pulchra*. C-type lectins are found in all xenacoelomorph species investigated [53,69]. Toll-like receptors (TLR) were described as lost in Xenacoelomorpha [51]. However, we find their characteristic domains [51,70] in the nemertodermatid *N. westbladi*, which was not included in the previous study. NOD-like receptors (NLRs) features [59] are present in all acoels investigated apart from *Praesagittifera naikaiensis*, but are absent from non-acoel Xenacoelomorpha. We find sequences from all species, however, containing NACHT domains without Leucin-rich repeats.

As signalling pathways, we examined the Toll pathway and the complement system. Three activation pathways for the complement system are known, all converging to the cleavage, and consequent activation, of C3 [53,71,72]. We therefore focused on C3, along with components of the alternative pathway, the most robustly conserved across metazoans [53,73–75] (Fig. S1B). We find xenacoelomorph predicted proteins containing α2-macroglobulin domains, but not the other domains of metazoans’ C3 [53,69,76–79] (Fig. S1B); besides, they are more closely related to other genes of the alpha-2-macroglobulin family than to C3 (Fig. S2) [79]. Factor B domain combination is not present in predicted proteins from the investigated xenacoelomorph species [53,77,80–83]. Sequences with the domain pattern of complement receptors 1 and 2 [53,80,84] are found only in the non-acoel xenacoelomorphs *X. bocki, N. westbladi,* and *Meara stichopi,* and they cluster together with metazoan CR1/2 (Fig. S1B, S3). The Toll pathway comprises the receptors TLRs, three signalling mediators–MyD88, Tube/IRAK4, Pelle/IRAK1–, and the transcription factor NFκB [53,85–90] (Fig. S1C). We find predicted proteins with both MyD88 domains [53,89,91] only in the non-acoel xenacoelomorphs *X. bocki, M. stichopi*, and *N. westbladi.* However, when running a phylogenetic analysis, these sequences do not cluster with other metazoans’ MyD88, but with human TIRAP, though with low support values (Fig. S4). The domains characterizing IRAK4/Tube and IRAK1/Pelle—Interleukin-1 receptor-associated kinases (IRAKs)—are not featured together in any predicted proteins from the investigated xenacoelomorphs [53,87]. While we find sequences from all species containing Rel/Nuclear Factor-κB (NFκB) domains [53,92,93], according to the phylogenetic analysis only *M. stichopi* sequences can be considered NFκB (Fig. S5). *Xenoturbella* and nemertodermatids also contain sequences more closely related to Rel proteins, and all the investigated species possess sequences clustering within the NFAT (Nuclear Factor of Activated T-cells) family (Fig. S5) [93].

Summarising, PRRs are mostly conserved in acoels, with the exception of TLRs. Signalling pathway components of the innate immune system (toll pathway and complement system) are partially absent in non-acoel xenacoelomorphs and completely absent in acoels.

### Mortality upon immune challenge increases in photosymbiotic acoels, not in aposymbiotic ones

We then investigated the link between photosymbiosis and immunity with *in vivo* immune challenges of photosymbiotic and non-photosymbiotic (*i.e.* aposymbiotic) acoels. Our hosts of choice are *Convolutriloba macropyga* photosymbiotic adults and aposymbiotic juveniles [37], as well as the aposymbiotic species *Hofstenia miamia* [94]. Since no pathogens of acoels are yet known, the Gram-negative bacterium *Vibrio coralliilyticus* was selected as immune agent [95]; *V. coralliilyticus* is a pathogen to a variety of marine organisms, such as corals, sea anemone, fish, bivalves, crustaceans, and sea urchin [95–103]. *V. coralliilyticus* has also been associated with disruption of symbiosis in corals [101,104–108], although not in sea anemone [103].

The animals were exposed for two days to a low or high bacterial load—10^5^ and 10^6^ CFUs (colony forming units), respectively. *C. macropyga* adult mortality is affected by bacterial dose and by batch (Fig. 2A, Table S2) The mortality is significantly higher for high-bacterial-load samples than for controls or low-bacterial-load samples (HR = 4.65 ± 0.21, Bonferroni-adjusted p<0.0001 for both post hoc contrasts), as well as for low-bacterial-load samples compared to controls, although with lower hazard and lower significance value (HR = 2.23 ± 0.23, Bonferroni-adjusted p=0.0012). To confirm the active role of *V. coralliilyticus* in causing *C. macropyga* mortality, we also performed immune challenges with heat-inactivated bacteria. A high-dose of heat-inactivated bacteria doesn’t increase mortality compared to controls (Fig. S6A). Exposure to the Gram-positive *Priestia megaterium* [109,110] was also tested and the mortality increase is even starker at a high bacterial load, yet none at a low bacterial load (Fig S6B, Table S2).

**Fig. 2.**
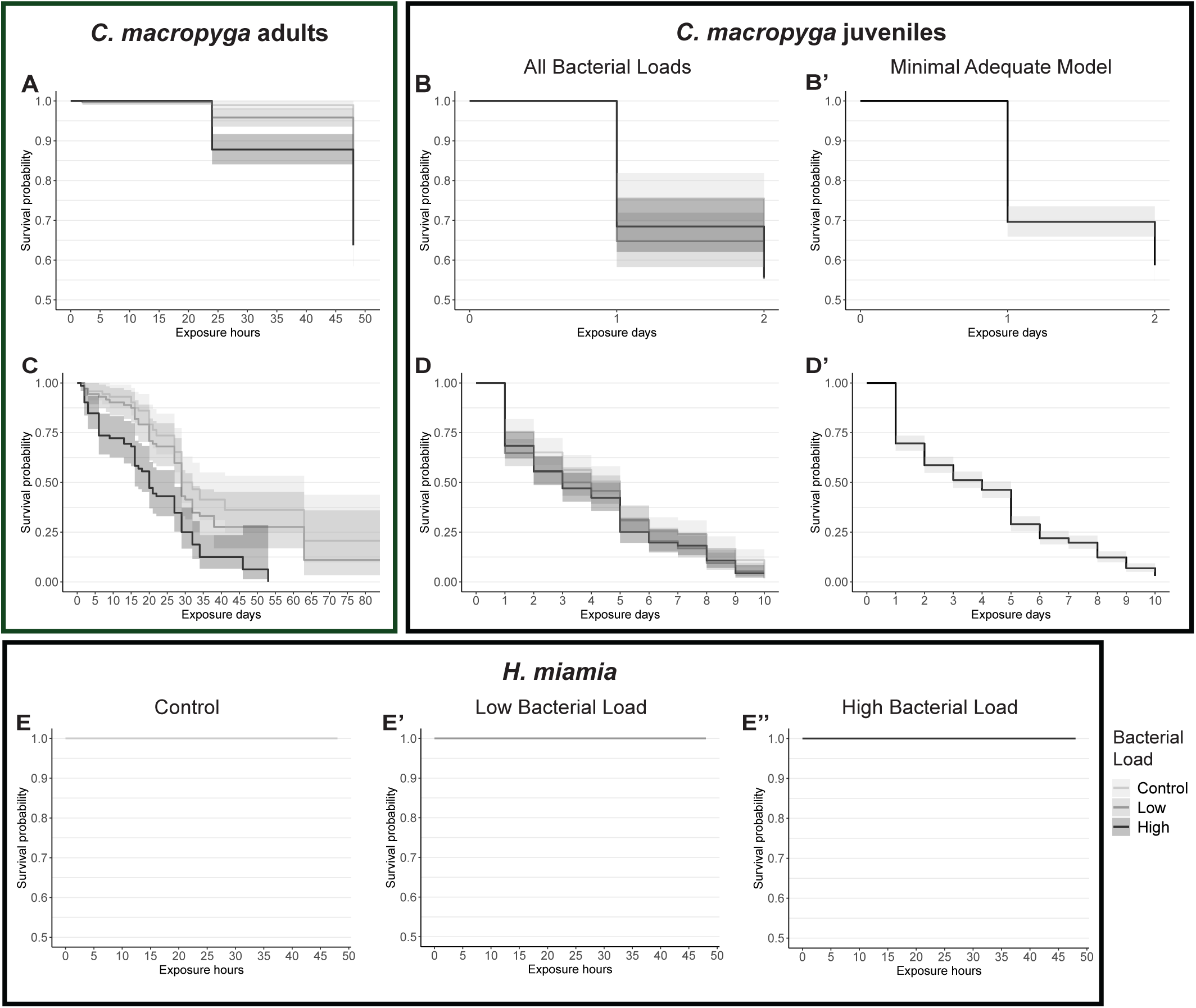
Survival curves of acoels upon *Vibrio coralliilyticus* exposure. Survival curves with 95% confidence intervals for immune challenges with *V. coralliilyticus.* Data were fitted to mixed effects Cox proportional hazard models, full statistics in Table S2. Two days of bacterial exposures: **A** *C. macropyga* adults (minimal adequate model Surv(last.obs, censored) ∼ Bacterial.Load + (1 | batch), χ^2^ = 59.742, p = 1.065e-13, number of replicates = 4, n = 863); **B-B’** *C. macropyga* juveniles (minimal adequate model Surv(last.obs,censored) ∼ 1, number of replicates = 4, n = 569); **E-E’** *H. miamia* adults (number of replicates = 5, n = 154). Long-term exposures: **C** *C. macropyga* adults (minimal adequate model Surv(last.obs, censored) ∼ Bacterial.Load + (1 | batch), χ^2^ = 8.646, p = 6.02e-07, number of replicates = 3, n = 216); **D,D’** *C. macropyga* juveniles (minimal adequate model Surv(last.obs,censored) ∼ 1, number of replicates = 4, n = 569). For *C. macropyga* juveniles, graphs representing the survival for all bacterial loads (**B,D**) are added alongside the ones showing the minimal adequate model (**B’,D’).** In (**B’,D’**) the colour of the curve is not meaningful, since survival is independent of bacterial load. Note that the y axis lower limit is 0.5 for the two-days graphs.

The mortality of *C. macropyga* juveniles is not affected by bacterial load nor by batch (Fig.2B-B’). Immune challenged *H. miamia* survived regardless of bacterial load, except one animal in the high dose sample (Fig. 2E-E’’). Besides, all individuals (n = 25) exposed to an even higher dose of 2.5·10^7^ CFUs also survived. In some wells, despite unaffected animal morphology, spheres of tissue could be observed in solution, and in one individual a sphere appeared from the posterior end of the body and another one was expelled through the mouth (video S1).

We also carried out longer immune challenge assays in *C. macropyga* adults and juveniles, to check for a delayed effect on survival. They were monitored for 10 days (juveniles), 1 month (2 batches of adults) or 3 months (1 batch of adults). In a similar way to the two-day assays, the bacterial load affected survival for adults and not for juveniles (Fig. 2C,D’).

To summarize, survival in aposymbiotic *H. miamia* and *C. macropyga* juveniles is seemingly not impacted by exposure to *V. coralliilyticus*, while mortality increases in symbiotic *C. macropyga* adults.

### Bacterial distribution within immune challenged *Convolutriloba macropyga*

To confirm infection of *C. macropyga* upon exposure, we performed *in situ* hybridisation against *Vibrio coralliilyticus* 16S rRNA on individuals exposed 2 days to a low bacterial dose (Fig. 3A). A variety of patterns can be observed, from individuals showing no signal at all (6 out of 34) to almost ubiquitous signal (14 out of 34), with multiple instances of signal localized in the digestive system area. In half of the control samples (8 out of 17), bacterial 16S rRNA is also observed around the anterior region of the digestive system (Fig. 3B). While in most of the other samples no signal can be found, in two of them bacterial RNA is present in the whole sample. In individuals exposed to *V. coralliilyticus* for shorter times, we see similar patterns, with signal for *V. coralliilyticus* 16S both in exposed individuals and in controls (Fig. S7). It can also be found in correspondence to the anterior nerve cords in some samples and in asexual reproductive buds. To confirm the presence of 16S rRNA from *V. coralliilyticus* in *C. macropyga*, we checked for its amplification by PCR from *C. macropyga* cDNA: a band can be seen for exposed samples, control ones, and for cDNA synthesized before the acquisition of *V. coralliilyticus* cultures (Fig. S8). The probe itself does not yield any hits when BLASTed against *C. macropyga* transcriptome, and the primers do not amplify anything *in silico* from *C. macropyga* transcriptome (see Methods). Therefore, it is reasonable to consider the *in situ* hybridisation signal as coming from 16S rRNA of *V. coralliilyticus* or a related bacterial species.

**Fig. 3.**
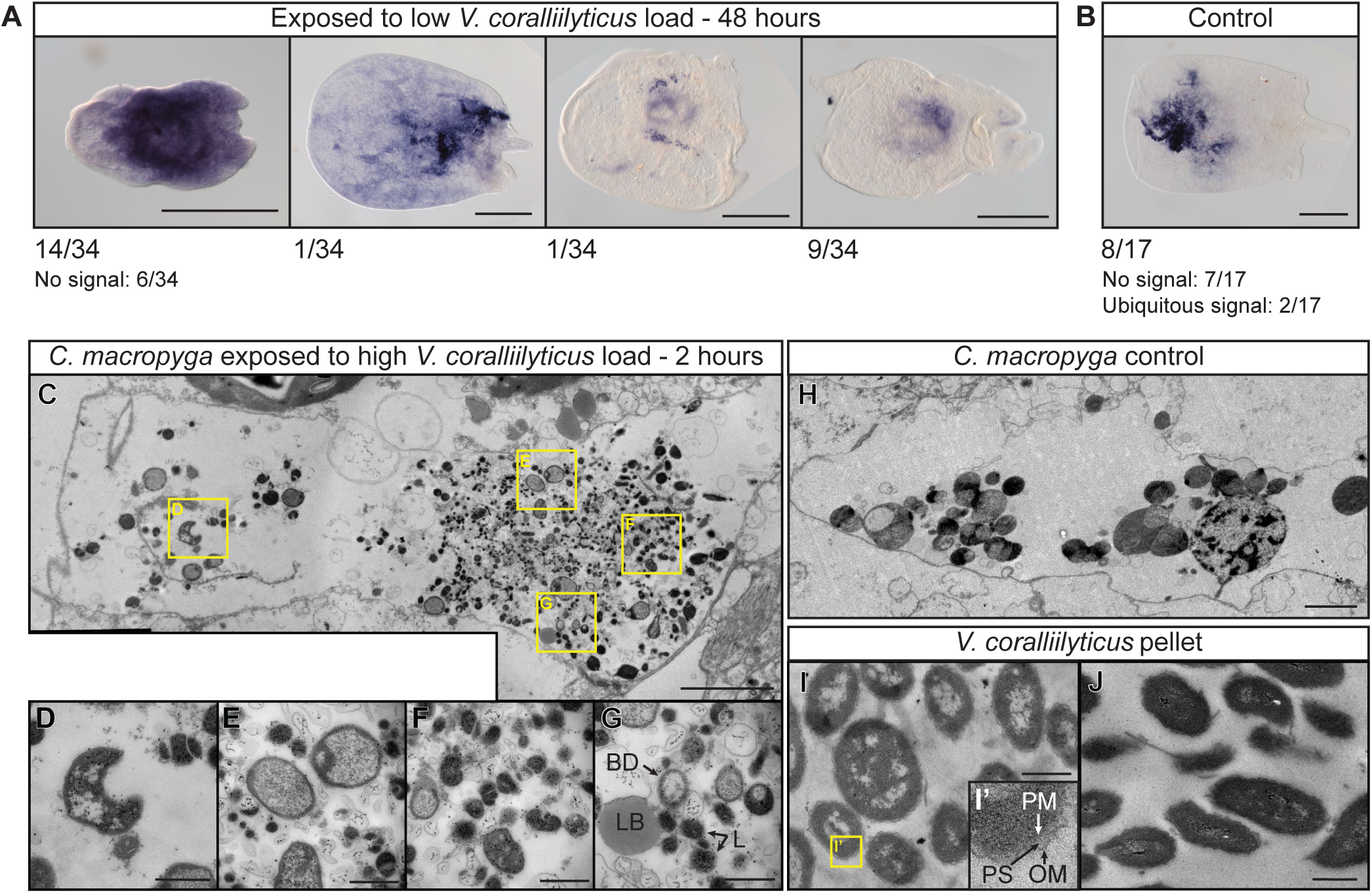
Presence of *Vibrio* in immune challenged *Convolutriloba macropyga*. RNA *in situ* hybridisation against *V. coralliilyticus* 16S in *C. macropyga* exposed for 48 hours to (**A**) a low load of *V. coralliilyticus* and (**B**) resuspended marine broth (control). Numbers indicate the ratio of individuals with the pattern shown above; dorsal view, anterior is facing left. Transmission Electron Microscopy images of phagocytosis in the digestive parenchyma in *C. macropyga,* cross section at the level of the mouth: **C-G** exposed to a high load of *V. coralliilyticus* for 2 hours, containing degraded bacteria; **H** in 2h-exposure control *C. macropyga*, showing physiological phagocytosis; **I,J** cultured and pelleted *V. coralliilyticus*. Yellow squares magnifications: **D** abnormal bacterial shape and absence of periplasmic space and outer membrane typical of gram-negative bacteria; **E** abnormal pale granulated cytoplasm of the pathogens; **F** numerous osmiophilic lysosomes; **G** bacterial debris (empty cell walls) (BD), lipid or lipofuscin body (LB), lysosomes (L) [111,112]; **I’** hallmarks of gram-negative bacteria: outer membrane (OM), a well-defined periplasmic space (PS), plasma membrane (PM) [113]. Scale bars are 0.5 mm in (A,B), 3 µm in (C,H), and 0.5 µm in (D-G,I,J).

To detect *V. coralliilyticus* in *C. macropyga* infected tissues and the host response we performed Transmission Electron Microscopical (TEM) on the brain, the body wall and internal parenchyma in the digestive system area (*i.e.* the areas showing signal for *V. coralliilyticus* 16S rRNA, Fig. 3A,B). We could not find any bacteria in individuals exposed for 2 days to a low (n=2) or a high load (n=2) of *V. coralliilyticus* or to control medium (n= 2). We therefore imaged individuals exposed for a shorter period: 1 hour of exposure (n=2) and 2 hours of exposure (n=2). Potential pathogenic bacteria at various levels of degradation were found in the digestive system of a 2-hours exposure individual (Fig. 3C-G). Signs of phagocytosis—such as lysosomes, phagolysosomes, lipid or lipofuscin bodies (Fig. 3F,G)—are found also in the digestive parenchyma of control samples (Fig. 3H). Some structures, however, resemble free-living *V. coralliilyticus,* but shrunk and with abnormal features (compare Fig. 3D,E to I,J). Similar characteristics are observed in *V. coralliilyticus* within infected corals [104,106], although in our samples a wider degree of variation can be seen.

### Symbiotic algae number decreases upon immune challenge in *Convolutriloba macropyga*

Disruption of the symbiosis between corals and algae is a known response to environmental stresses and infection by some bacteria, among which *Vibrio coralliilyticus* [114]. To investigate this in *C. macropyga* immune challenged with *V. coralliilyticus*, we first looked for whitening of the whole animal or parts of it, as observed in corals [104,106,115]. No white areas can be distinguished at 2 days of exposure in fixed and mounted individuals (Fig. 4A-C). At 14 days of exposure, when bleaching is visible in *V. coralliilyticus*-infected corals [104,106], still no pigmentation change can be detected in *C. macropyga* (Fig. S9). We then quantified potential symbiont loss at the tissue level: algae chlorophyll autofluorescence and stained animal cell nuclei were imaged by confocal microscopy in a relatively flat area between the eyespots and mouth (Fig. 4D-D’’), then automatically detected and counted. At 2 days of exposure, the ratio between algal cells and animal cells is affected by bacterial load (Fig. 4E, Table S3). Post hoc Tukey HSD tests (Fig. 4E) show that individuals exposed to a high dose of *V. coralliilyticus* have 1.7 less algae per animal cell than controls, while comparisons of low-dose samples to the other two conditions are not significant. While the algae per animal cell ratio is not affected by animal size (table S3), exposure to *V. coralliilyticus* has an effect on animal size itself (Fig. 4F). Both a high dose and a low dose of *V. coralliilyticus* decrease animal size, measured as length, though the effect is starker and more robustly significant for the high dose (Fig. 4F).

**Fig. 4.**
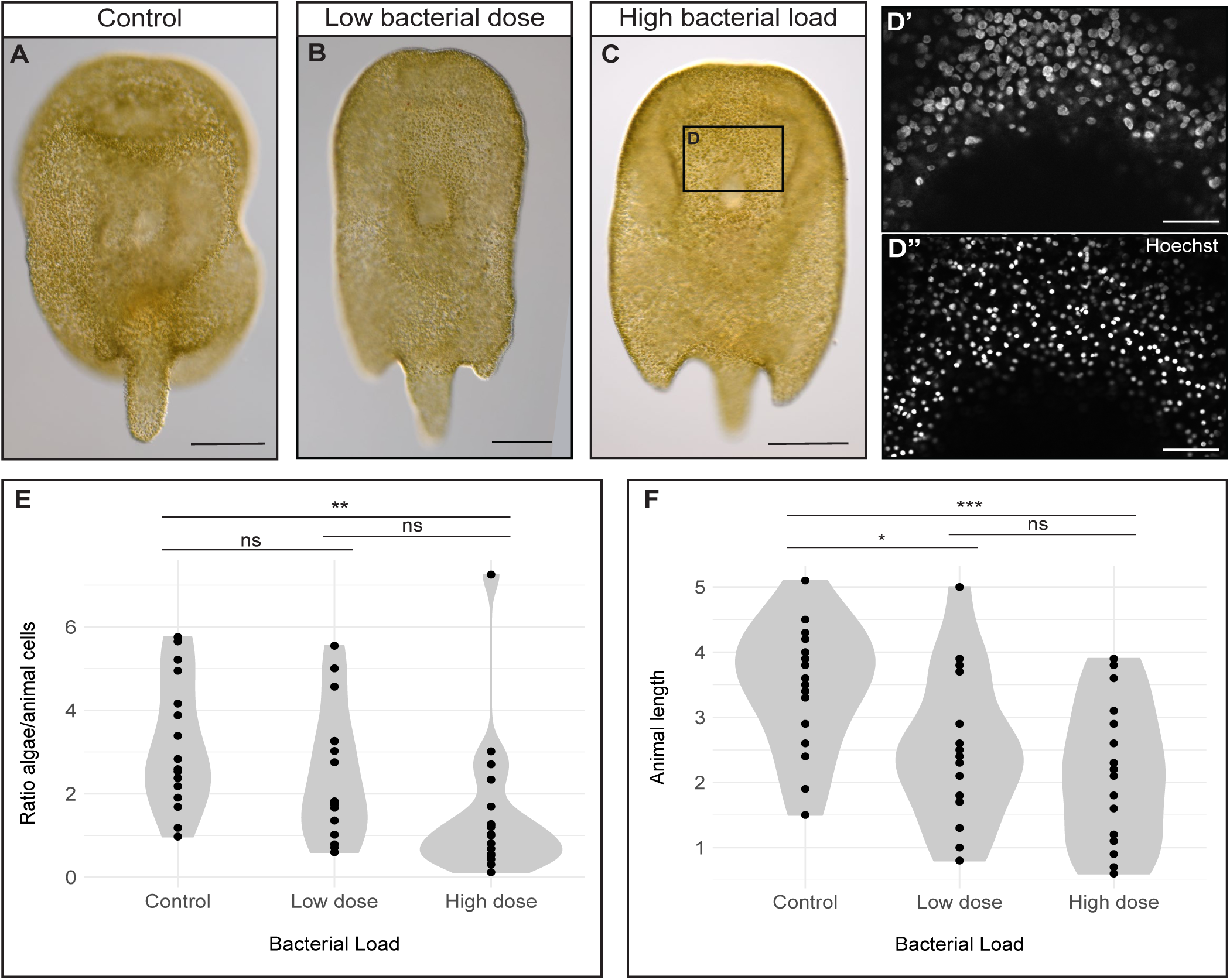
Dysbiosis in *C. macropyga* upon 2-days immune challenge with *V. coralliilyticus*. **A-C** DIC-images of *C. macropyga* after 2-days immune challenges with *V. coralliilyticus*; ventral view, anterior facing up. **D’-D’’** Chlorophyll autofluorescence and Hoechst 33342 staining in the area corresponding to (**D**), as an example of the data used to produce the plots in (**E**). **E** Violin plots of the ratio of algal cells to animal cells in immune challenged animals. Minimal adequate model: Ratio.algae.hoechst ∼ Bacterial.Load, number of replicates = 2, sample size = 52, F = 5.5069, p = 0.00696). Post hoc Tukey HSD tests: high-dose – control t = −3.298, p = 0.00514; low-dose – control t = - 1.348, p = 0.37598; low-dose – high-dose t = −1.861, p = 0.16080. Detailed statistics in Table S3. **F** Violin plots of animal length in immune challenged animals. Length ∼ Bacterial.Load, Anova, F = 7.7059, p = 0.001231; post hoc Tukey HSD tests: high-dose – control t = −3.847, p < 0.001; low-dose – control t = −2.622, p = 0.0306; high-dose – low-dose t = −1.093, p = 0.5229. For both graphs, p values for Tukey HSD tests: ns: p > 0.05; *: p < 0.05; **: p < 0.01; **: p < 0.001. Scale bars are 250 µm in (**A,B**), 50 µm in (**D’-D’’**).

### Damaged tissue in immune challenged *Convolutriloba macropyga*

Immune challenged *Convolutriloba macropyga* were considered dead when not moving and showing loss of anatomical integrity. They shrink and present structural deterioration, with pieces coming apart (Fig. 5D). The same, however, can be observed for the death of control individuals (Fig. 5B) and is therefore not specific to *Vibrio* pathogenesis.

**Fig. 5.**
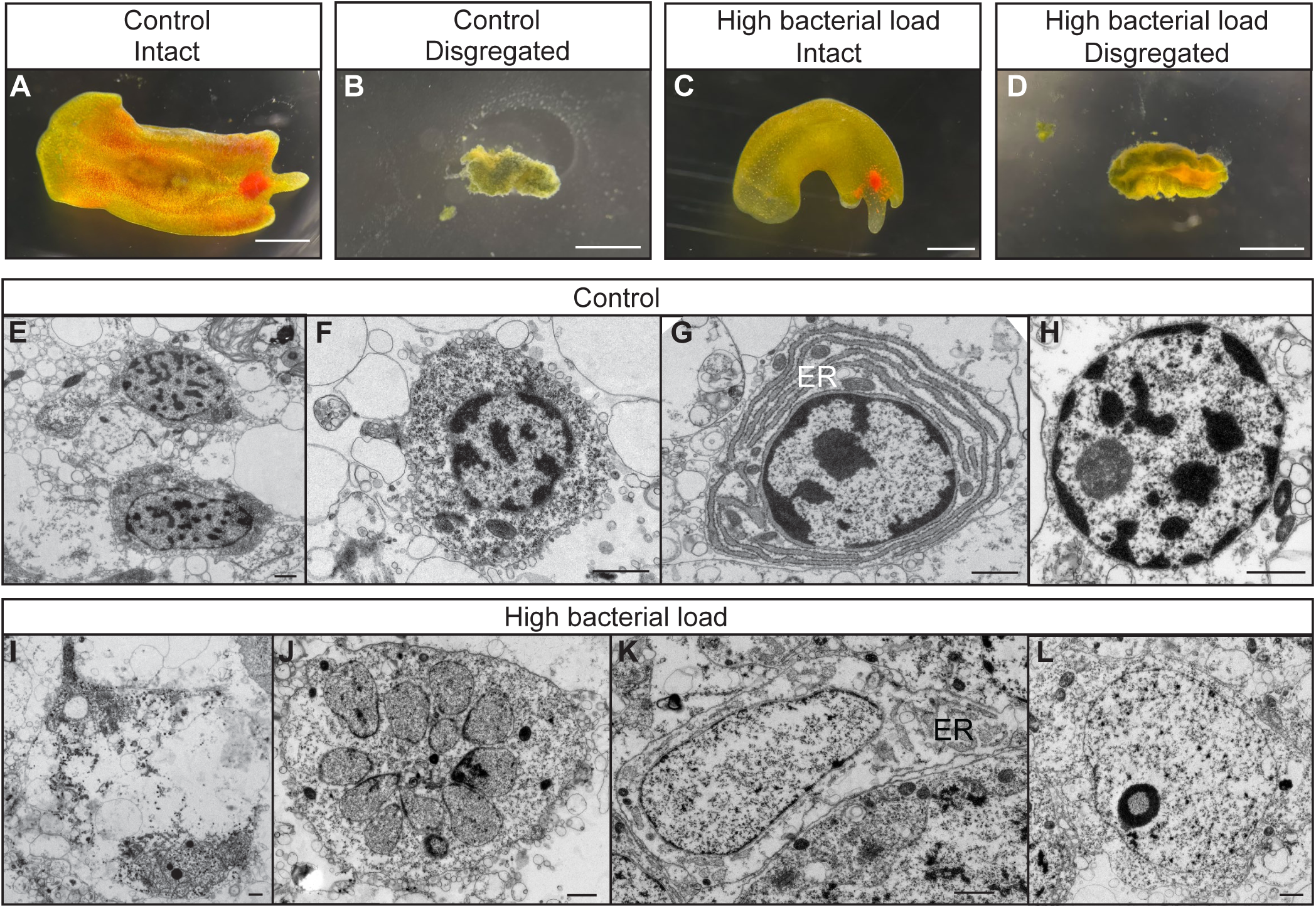
Damage in *C. macropyga* upon *V. coralliilyticus* exposure. *C. macropyga* at 2 days of exposure to *V. coralliilyticus*, imaged through a stereo microscope (**A-D**), or at ultrastructural level with Transmission Electron Microscopy (**E-L**). Features of parenchymal cells in control samples: **E,F** granular cytoplasm and nuclei with heterochromatin, **G** endoplasmic reticulum (ER), **H** nucleus with visible nucleolus. Samples exposed for 2 days to a high load of *V. coralliilyticus:* **I** tissue degradation and degenerate organelles; **J** fragmented nucleus; **K** lysis of chromatin and swollen ER; **L** damaged nucleolus. Scale bars are 1 mm in (**A-D**), 1 µm in (**E-L**).

We searched for changes at the ultrastructural level with transmission electron microscopy in immune-challenged individuals that looked intact and motile. Tissue and cell damage is observed in *C. macropyga* exposed for 2 days to a high bacterial dose (Fig. 5I-L, n=2), when compared to controls (Fig. 5E-H, n=2). Tissue degradation and cell debris are present in the parenchyma (Fig. 5I) [cf. 106]. At a subcellular level, one can see swelling of the endoplasmic reticulum (Fig. 5K) [cf. 114], and segregation of nucleolus components (Fig. 5L) [cf. 117]. Fragmented nuclei are also present (Fig. 5J), although the lack of chromatin condensation and membrane blebbing rules out apoptosis [118,119].

### Asexual reproduction is not affected by *Vibrio* exposure

Reproduction contributes to individual fitness alongside survival, and the impact of infections on it can be antipodal [120]. In fact, on one hand, pathogens have been shown to decrease fecundity, likely due to energy constraints [121–123]. On the other hand, organisms sometimes allocate more energy to reproduction when their survival is threatened, a strategy known as terminal investment in reproduction or fecundity compensation [120,124–128]. We therefore measured fecundity by monitoring the number of asexual progeny released per day upon 2-days immune challenges. Asexual progeny numbers are not affected by bacterial load. They are however affected by presence of an asexual bud prior to bacterial exposure and, in the first day of exposure, by batch (generalized linear mixed model, minimal adequate model for day 1: progeny.released ∼ bud + (1 | batch), χ^2^= 24.684, p = 6.752e-07, number of replicates = 4, sample size = 813; day 2: progeny.released ∼ bud, χ^2^= 18.512, p = 1.688e-05, number of replicates = 4, sample size = 669; detailed statistics in Table S4). Therefore, no changes in reproduction occur upon *V. coralliilyticus* exposure, neither an increase nor a decrease.

### PRRs are expressed around the digestive system, nervous system, and in reproductive structures

To characterize acoel molecular response to infection, we first focused on *C. macropyga* Pattern Recognition Receptors (PRRs) identified as detailed above. Expression patterns for each PRR gene vary widely between individuals, both after exposure to a low dose of *V. coralliilyticus* and in controls (Fig. 6). They are not expressed in some samples at all, and they are expressed in the whole animal in some other samples. The remaining cases can be described as follows. C-lectin gene expression localises around the mouth opening or more broadly in the digestive system for control samples (Fig. 6A-C). Upon bacterial exposure, the changes in expression vary by gene, with one C-lectin widening its expression domain (Fig. 6A, also corresponding to Scavenger Receptor E domain structure, Fig. S1), another showing no change (Fig. 6B), and a third not being expressed at all in the exposed samples (Fig. 6C). The NOD-like receptor gene (NLR, Fig. 6D) is expressed in the anterior part of the digestive system and in the nervous system in controls. It either widens or narrows its expression pattern in immune challenged animals. Scavenger Receptors Type B (SR-B, Fig. 6E-F) are expressed throughout the body, with increased intensity around the digestive system in controls. Other two domains of expression on the sides of the digestive system could correspond to two nerve cords or to the female gonads. The pattern is less clear-cut for exposed individuals and there seems to be stronger expression in the anterior region of the digestive system, as well as in the nervous system. Even if a clear pattern cannot be derived for all PRRs, we can identify some areas of increased immune receptor presence in acoels (Fig. 6H). Asexual reproduction buds at the posterior end of the animal body, when present, express the immune receptors. Other putative hot spots of immune recognition are the areas around the mouth and around the digestive system, the anterior nervous system, the female gonads – when present –, and a faint domain on the flanks of the animals, which, to our knowledge, does not correspond to any described morphological structure in acoels.

**Fig. 6.**
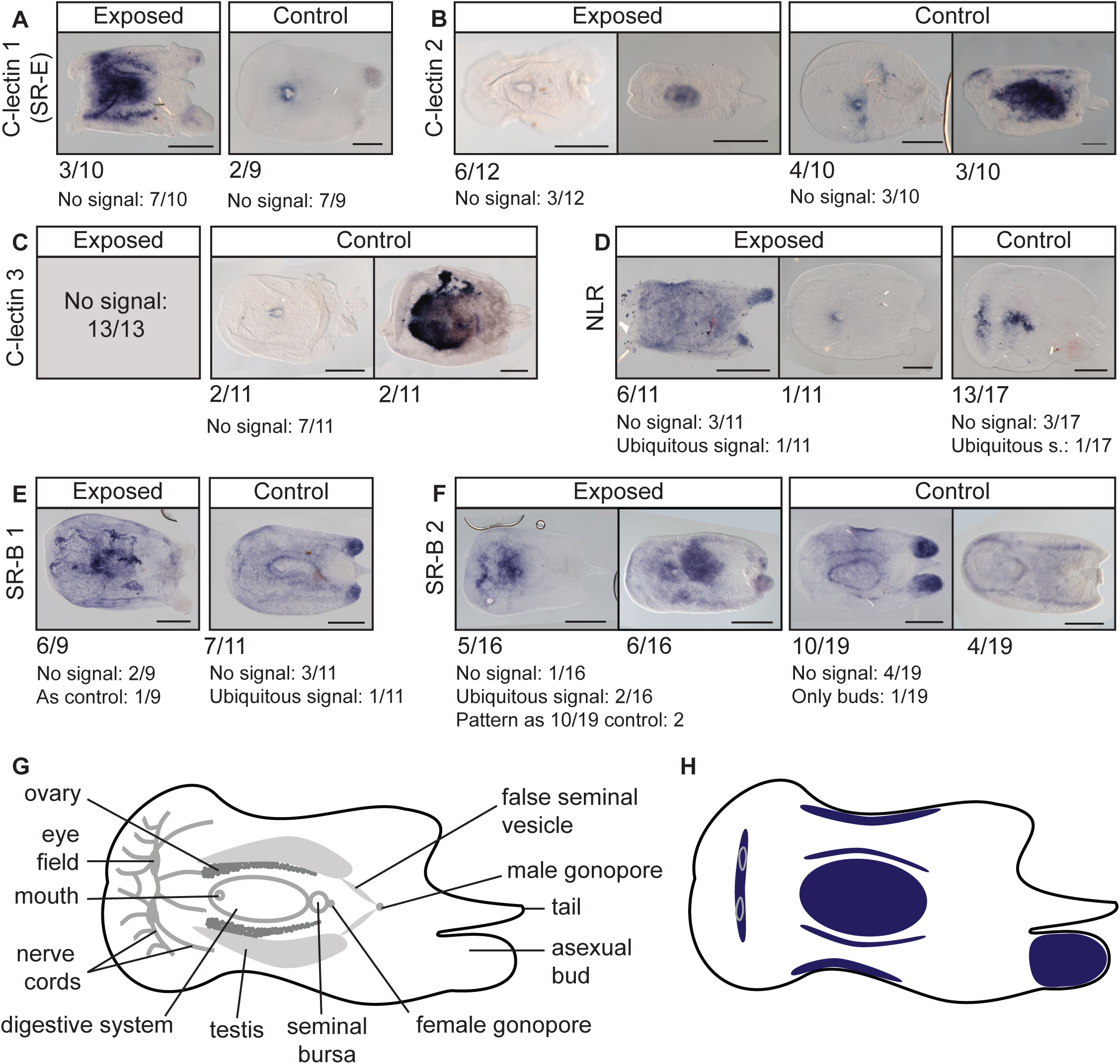
Expression patterns of *Convolutriloba macropyga* PRRs upon immune challenge. **A-F** RNA *in situ* hybridisation against *C. macropyga* PRRs in immune challenged *C. macropyga* adults (exposed for 48 hours to low *V. coralliilyticus* load or control medium). Numbers indicate the ratio of individuals with the pattern above; dorsal view, anterior facing left; scale bars are 0.5 mm. **G** Morphology of *C. macropyga* after Shannon & Achatz [37]. **H** Schematic of PRR expression domains.

### Many genes differentially expressed upon immune challenge are acoel-specific or not characterized

Given the lack of canonical immune signalling pathways, we looked for genes involved in the response to *Vibrio* by comparing gene expression levels in *C. macropyga* adults challenged with a low *V. coralliilyticus* load to controls at 2 days of exposure. Principal component analysis of count data after variance stabilizing transformation shows that the samples cluster by date of infection (batch) and not by condition, *i.e.* control or exposed (Fig. 7A). Thus, most of the differences between samples are due to batch, not to bacterial exposure status. 29 genes are differentially expressed upon exposure to *V. coralliilyticus* (8 upregulated, 21 downregulated; FDR-adjusted p < 0.01; Fig. 7B,C, Table S5). When searched against the NCBI non-redundant protein sequence database, half of them (14/29) retrieve uncharacterized proteins or no hits at all. Of those, two match uncharacterized proteins in multiple eukaryotic species, two have no correspondence in xenacoelomorph predicted proteomes, 8 only in *C. macropyga*, and two in other acoels too—but not in non-acoel xenacoelomorphs (Fig. 7C, Table S5). Four differentially expressed genes match bacterial sequences or conserved domains, though none of them corresponds to *Vibrio* species: hits for the two downregulated sequences are mostly uncharacterized bacterial sequences, while the two upregulated sequences seem related to virulence factors (one as WXG100 family [129]; one with domains corresponding to MAC/Perforin, Hint and YeeP GTPase [130]). Hits for one downregulated sequence are secreted RxLR effector protein 161-like for the soft coral Xenia sp. Carnegie-2017 (automatically predicted); though these are animal sequences, RxLR effector proteins are usually avirulence factors from pathogenic oomycetes [131]. One sequence matches only an unnamed protein from the green alga *Closterium* sp. and could therefore correspond to an algal sequence. Hits for all other sequences are metazoan genes: one upregulated serine dehydratase, one upregulated transient potential cation channel, 3 downregulated serine proteases, one downregulated dynein regulatory complex subunit, one downregulated adhesion protein, one downregulated carbonic anhydrase, and one downregulated C-lectin domain-containing protein— which does not align to the ones retrieved in our initial gene search. While the latter is the only canonical immune-related gene, serine proteases can also be involved in immunity [132].

**Fig. 7.**
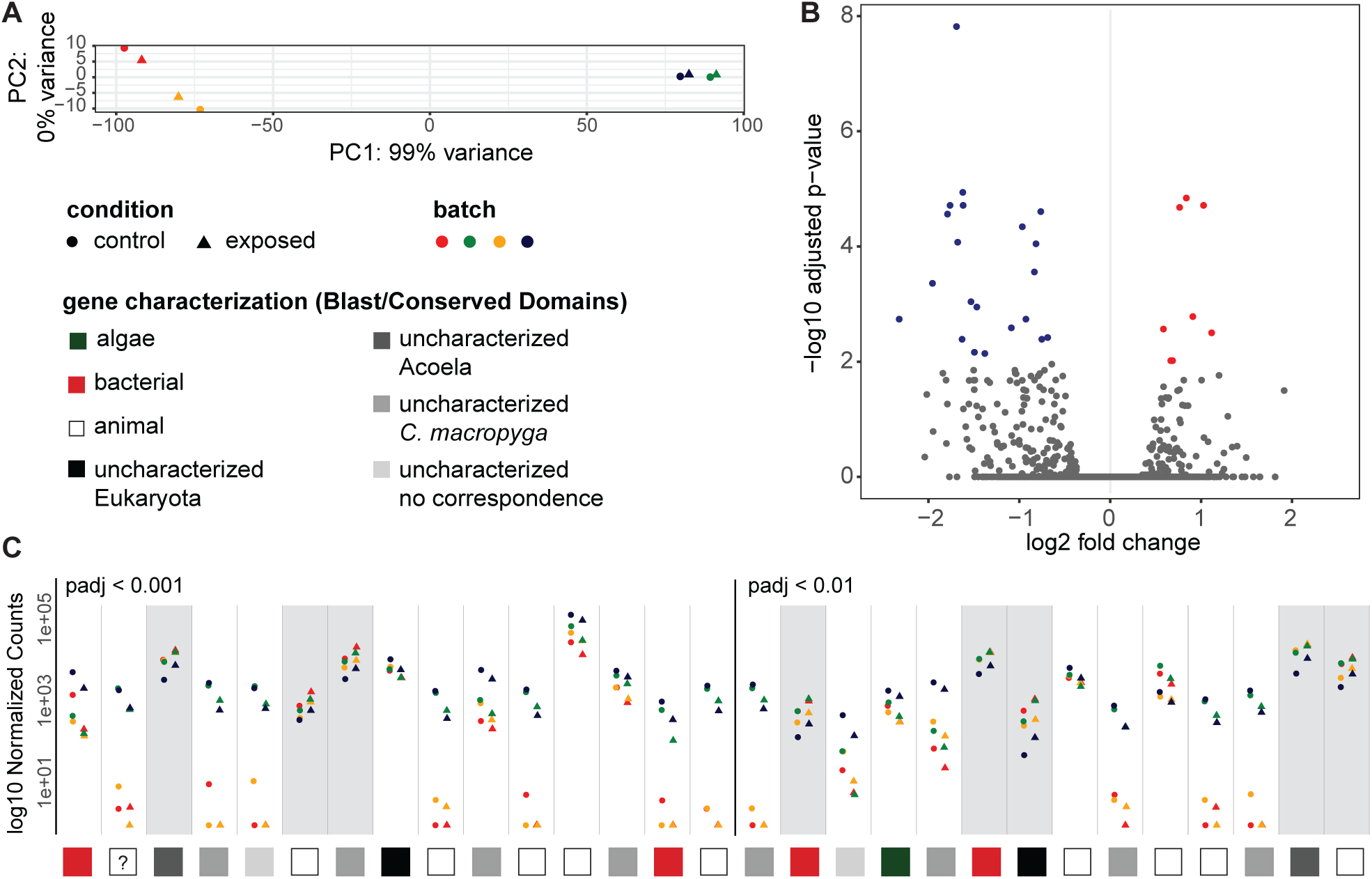
Differentially expressed genes upon immune challenge. **A** Principal Component Analysis of transformed count data. **B** Volcano plot of differentially expressed genes (DEGs) between control and exposed samples. Blue dots represent downregulated genes, red upregulated, grey genes below significance threshold (padj<0.01). **C** Normalized counts for significantly DEGs by batch, ordered by increasing adjusted p-value. Genes with grey background are upregulated, with white background downregulated. Data point shapes and colours are common to (**A**) and (**C**). Squares in (**C**) show the results of gene characterization by BLAST and Conserved Domain Search, detailed in Table S5. The question mark identifies the sequence corresponding to *Xenia* sp. RxLR effector protein.

## Discussion

### Photosymbiosis in Xenacoelomorpha evolved after the loss of metazoan-conserved immune signalling pathways

We analysed the literature on the presence and type of photosymbionts in xenacoelomorphs (Table S1), mapped them on the phylogeny (Fig. 1A), and consequently confirmed that photosymbiosis has likely evolved at least twice in acoels, always within Convolutidae [38]. One group includes our focus species *Convolutriloba macropyga* and the better known *Symsagittifera roscoffensis*; photosymbionts in this group are always green algae, with the exception of *Convoluta convolutae*, which bears diatoms as endosymbionts [133, Table S1]. In the other group, comprising *Waminoa* and *Amphiscolops*, the endosymbionts are dinoflagellates; some *Amphiscolops* species can simultaneously host dinoflagellates and green algae [38,134] (Table S1). In Japan, Riewluang & Wakeman [135] recently found a novel group of photosymbiotic acoels outside of Convolutidae, as a sister group to Mecynostomidae. These species were not included in our sources for phylogeny [45,46] and are therefore not shown in our analysis, but a third independent occurrence of photosymbiosis remains possible.

Among the immune genes usually conserved in metazoans, there are some Pattern Recognition Receptors (PRRs) and genes belonging to signalling pathways, such as the complement system and the Toll pathway [53] (Fig. 1B). We searched for predicted proteins with a domain structure corresponding to known PRRs in Xenacoelomorpha and found C-lectins, as well as Scavenger Receptors of class B and E. Toll-like receptors (TLRs) are considered lost in xenacoelomorphs [51]; we only recover them in the nemertodermatid *Nemertoderma westbladi*, a species not included in previous analyses [51]. A more in-depth investigation would be required to understand if *N. westbladi* sequences are homologous to other metazoan TLRs or if they have evolved *de novo,* for example from TIR-only or LRR-only containing proteins. NOD-like receptors (NLRs) are present in all investigated acoels, but *Praesagittifera naikaiensis*; they could not be found in *Xenoturbella* and nemertodermatids.

As for genes belonging to the complement system, they are progressively lost in xenacoelomorphs and completely missing from acoels. In fact, C3—the pathway central component—and factor B are absent from xenacoelomorphs and complement receptors 1/2 are only present in non-acoel xenacoelomorphs (Fig. 1B). The other signalling pathway we investigated, the Toll pathway, also seems completely lost in acoels and only traces can be found in other xenacoelomorphs (Fig. 1B). The activators of the pathway, TLRs, are only recovered in one of the two nemertodermatid species investigated, as discussed above. The transcription factor usually activated by the signalling cascade, NFκB, is only convincingly detected in the other nemertodermatid species, *Meara stichopi.* Predicted proteins with MyD88 domain structure (TIR and DD) are only found in non-acoel xenacoelomorphs. Upon phylogenetic analyses, these sequences could be more closely related to human TIRAP (TIR-domain containing adaptor protein) than to other metazoan MyD88s (Fig. S4). TIRAP is also involved in the Toll signalling pathway, but is thought to be present only in chordates [136,137]. Therefore, if a MyD88-like adaptor protein exists in xenacoelomorphs, it cannot be found in acoels, and it diverged consistently from other metazoans’ MyD88. Our data thus suggests that a canonical metazoan Toll signalling pathway is lost in acoels and possibly in other xenacoelomorphs, too.

The PRRs we investigated are known or suspected to interact with photosymbiotic endosymbionts in cnidarians [1,19,27], and in gastropod molluscs [58,138]. Their conservation in acoels, with the exception of TLRs, prompts future investigations into their role in the interaction between acoels and their photosynthetic endosymbionts, too. The complement system and toll pathway are negatively associated with symbiosis in cnidarians: they are either downregulated in symbiotic animals or tissues, or upregulated during bleaching and loss of symbionts [27,62,82,139,140]. The loss of these signalling pathways precedes the evolution of photosymbiosis in acoels. It could have favoured the occurrence of photosymbiosis, especially considering that the endosymbionts are not confined to a specific tissue, as in cnidarians, but dispersed throughout the animal’s body instead [37].

### Coral-bleaching agent *Vibrio coralliilyticus* causes mortality and dysbiosis in photosymbiotic acoels

In corals, *Vibrio coralliilyticus* is an opportunistic pathogen, associated to bleaching, tissue lysis, and mortality [25,104,106,115]. It also causes mortality in a variety of marine animals, from bivalves and crustaceans to sea urchin and fish [96,98–100,102]. Our data shows that *V. coralliilyticus* has the potential to be virulent to photosymbiotic acoels, too: (i) Mortality in *Convolutriloba macropyga* adults increases upon exposure to *V. coralliilyticus* (Fig. 2A,C); (ii) *V. coralliilyticus* 16S rRNA is present in immune challenged animals, but also in controls (Fig. 3A,B, Fig. S8); (ii) heat-inactivated bacteria do not decrease survival (Fig. S6A), confirming an active role of bacteria in causing mortality. Tissue degradation (Fig. 5) also resemble the effects of *V. coralliilyticus* on corals [95,104,106] and sea anemones [103]. The presence of *Vibrio* rRNA in control samples suggests that these bacteria are common symbionts of *C. macropyga,* and that they can become virulent at higher loads. This contrasts with the fact that we haven’t observed any alive bacteria by Transmission Electron Microscopy, although this method doesn’t allow a comprehensive study and bacteria could simply have been missing in the sections or individuals observed. We also do not know if these laboratory observations correspond to conditions in the wild; thus, *V. coralliilyticus* cannot be definitely labelled as an acoel pathogen [141]. Data on *V. coralliilyticus* geographical distribution are available [142,143], although *C. macropyga* localization in the wild is still unknown [37]. Moreover, *V. coralliilyticus* virulence in corals is only observed at higher temperatures [142], with mortality only happening above 25°C or 27°C and bleaching above 24°C [104,106]. As for sea anemone, mortality—but not dysbiosis—is reported in *Exaiptasia diaphana* exposed to *V. coralliilyticus* at 30°C and not at 25°C by one study [144] and at 22°C by another [103]. Therefore, our results for mortality and dysbiosis—obtained at 26°C—could also vary with temperature.

It should be noted that, in cnidarians exposed to *V. coralliilyticus,* tissue loss is detectable only after 10-12 days and death after 15-21 days [104,106]. In contrast, an increase in mortality of *C. macropyga* can already be detected at 2 days of bacterial exposure. Thus, these acoels could be used as an early warning system for coral diseases in tropical aquaria or coral reef monitoring, in a similar way to rosebushes for vineyards.

However, mortality does not increase when *V. coralliilyticus* exposure happens in the absence of photosymbiosis, *i.e.* in aposymbiotic *C. macropyga* juveniles and aposymbiotic acoel species *Hofstenia miamia* (Fig. 2, Table S2). Thus, the presence of photosynthetic endosymbionts seems linked to a higher susceptibility to pathogens. This could be due to (i) a weakened immune status needed to maintain symbiosis, as suggested for corals [21] or (ii) host mortality caused by a detrimental effect of *V. coralliilyticus* on the endosymbionts rather than on the host itself; damages to symbiotic dinoflagellates were also observed in corals infected by *Vibrio* [145,146].

To our knowledge, there is no study directly comparing mortality in symbiotic and aposymbiotic cnidarians upon *V. coralliilyticus* exposure. However, different strains of *Symbiodinium* endosymbionts correlate with differences in disease resistance [147], suggesting that photosymbionts can affect a host’s ability of facing infections. In the sea anemone *Exaiptasia diaphana,* survival upon infection with bacteria *Pseudomonas aeruginosa* or *Serratia marcescens* is higher in aposymbiotic individuals compared to symbiotic ones, but this effect is reversed in the presence of starvation [148]. In the scyphozoan *Cassiopea xamachana* challenged with *S. marcescens,* survival is also higher in aposymbiotic than in symbiotic polyps [149]. Given a similar effect of photosymbiosis on disease susceptibility in independent photosymbiotic systems, this may be due to intrinsic features of photosymbiosis.

Since we compared *V. coralliilyticus* effects on mortality in different species or ontogenetic stages, these factors could be more relevant than the presence of photosymbiosis. For example, a meta-analysis in bilaterians with separate sexes showed that increased mortality upon immune challenges is associated with adult, but not juvenile stages [150]. An alternative explanation for *H. miamia* higher survival upon immune challenge could be its stronger regeneration capacity: although, whole body regeneration in *H. miamia* and *Convolutriloba* species has been described as comparable [151], damaged *Convolutriloba* species release a toxin that can cause the death of the animal itself and surrounding individuals [37,40,41]. Since signs of tissue disruption upon bacterial exposure are seen both in *H. miamia* and *C. macropyga* (Fig. 5, video S1), *H. miamia* could be more efficient at repairing damage than *C. macropyga*.

The reduced number of algal symbionts per animal cell suggests that photosymbiosis breaks down upon *V. coralliilyticus* exposure (Fig. 4, Tables S3). While coral dysbiosis is usually clearly recognizable by white areas, in the scyphozoan *Cassiopea andromeda* elevated sea temperatures cause “invisible bleaching”, a decrease in symbiont density and chlorophyll activity without any noticeable pigmentation change [152]. A similar phenomenon could also be at play in sea anemones challenged with *V. coralliilyticus*, where no bleaching is visible [103]. The shorter length of *C. macropyga* exposed to *V. coralliilyticus* could be linked to a loss of nutrient apport from the symbionts. However, in *Convolutriloba retrogemma* exposed to elevated temperatures, a decrease in symbiont numbers was observed prior to mortality, without any changes in animal length [43]. Therefore, symbiont loss and mortality can happen without a size decrease in *Convolutriloba* species. This suggests that either responses to heat stress and immune challenges differ, or that shrinking in immune challenged individuals is not due to symbiosis disruption.

### Acoels are likely to employ novel molecular immune response mechanisms

So far there have been no studies on acoel immune responses to pathogens. Given the lack of usually highly conserved immune pathway genes (Fig. 1B), non-canonical immune mechanisms may play a role. We initially checked for a behavioural response to infections: terminal investment in reproduction. An increase in reproduction, in lieu of a costly immune response, could be linked to a reduced immune gene repertoire and immune response in pea aphids [153–155]. However, progeny release is unaffected in immune challenged *Convolutriloba macropyga*, so we rejected the hypothesis of terminal investment.

Transmission Electron Microscopy data in immune challenged *C. macropyga* show a degradation of the pathogens in the digestive parenchyma (Fig. 3C-G). *In situ* hybridisation also shows *Vibrio* rRNA and PRR (Pattern Recognition Receptor) expression inside or around the digestive system in most cases (Fig. 3A,B, Fig. S7, Fig. 6). The expulsion of a tissue ball by immune challenged *H. miamia* (Video S1) suggests the removal of digestive pathogens through the mouth. However, we did not observe a similar behaviour in *C. macropyga*, which could be linked to its less efficient response to *Vibrio*.

A molecular response to pathogen exposure also seems inconspicuous. First, *in situ* hybridisation shows that the PRRs identified in the predicted proteomes are expressed by the animal, but their expression domains vary greatly both in controls and in individuals exposed to *Vibrio* (Fig. 6). Nonetheless, we identified some regions that are more likely to be involved in the first stage of immune responses in acoels (Fig. 6H). The mouth and digestive system are likely the areas of first encounter and elimination of microbes, as discussed above; the nervous system and reproductive structures (gonads and asexual buds) would be essential areas to protect from pathogens. PRR expression in the digestive and nervous system has also been described for nemertean, nematode, and crustacean C-lectins [29,156,157], crustacean and mammalian SRs type B [156,158,159], and leech and mammalian NLRs [160–162]. *C. macropyga* flank regions expressing PRRs could correspond to a previously undescribed structure important for responses to microbes. Given the high variability in *V. coralliilyticus* 16S rRNA *in situ* hybridisation, the expression domains of PRRs could also be linked to the presence of bacteria, but we did not manage to perform double *in situ* hybridisation to test this. Secondly, our transcriptomic analysis indicates downregulation of one putative C-lectin, but no differential expression of other PRRs upon immune challenge (Table S5). Their expression levels could, however, peak early in response to pathogens, as shown in nemerteans and corals [53,163]. If this is the case, they would be already back to control levels (or decreased, as for the one C-lectin) at 48 hours of exposure, when our experiments were performed.

Moreover, in transcriptomic analysis, sample similarity correlates with batch rather than exposure status (Fig. 7A). This denotes a lack of substantial changes in gene expression upon *Vibrio* exposure, suggesting that molecular immune responses may be minor, at least at 48 hours of exposure. Upon differential gene expression analysis (Fig. 7C, Table S5), we find significant up- and downregulation of sequences belonging to non-Vibrio bacteria. This suggests an interaction between a potential pathogen and the bacterial community within *C. macropyga*, as extensively studied in human and model organisms [164,165]. The upregulated host transcripts we could characterize are serine dehydratase and transient receptor potential cation channel M. Serine dehydratase is an enzyme converting serine into pyruvate [166], the starting substrate for Krebs cycle [167]. Its upregulation could be linked to an increased energetic need following the reduction in nutrient-supplying endosymbionts. Carbonic anhydrase, which catalyses the conversion between CO_2_ and HCO_3_⁻, is important for photosynthesis and for photosymbiosis in anthozoans [168]. Its downregulation upon immune challenge could also be due to symbiosis breakdown. Transient receptor potential cation channels are involved in thermosensation and other sensory functions [169,170] and are expressed in putative sensory neurons in the acoel *Aphanostoma pulchra* [171]. Among the downregulated genes we find serine proteases: protein-cleaving enzymes with a variety of functions, which have been associated with several immune responses in insects and humans [132,172–174]. Downregulation of dynein regulatory complex subunit, a gene involved in ciliary function [175], and of adhesion molecules—related to trophinin and mucins [176], nidogens [177], and adhesion plaque protein [178]—could be linked to changes in cell motility. To summarize, changes in host gene expression levels seem to relate to potential immune responses, sensory functions, cell motility or adhesion, and the response to symbiosis breakdown. However, many of the genes differentially expressed upon immune challenge are not orthologues of any characterized gene and do not present known conserved domains (Fig. 7C, Table S5). Acoel-specific genes, as well as orthologues of still uncharacterized animal genes, seem to play a role in responses to bacterial exposure in acoels. Considering the unknown identity of these genes, the small significance of molecular responses, and the lack of changes in reproduction, novel immune strategies may be employed by acoels. Further investigation could uncover mechanisms relevant to human activities or even human health.

### Conclusion

Our study finds that Acoela, and to some degree the whole Xenacoelomorpha clade, lost immune pathway genes commonly conserved in animals. This loss preceded—and potentially favoured—the evolution of photosymbiosis, which happened at least twice independently in this clade. We show that exposure to the bacterium *Vibrio coralliilyticus* does not affect non-photosymbiotic acoels, but it increases mortality in the photosymbiotic acoel *Convolutriloba macropyga*. It also decreases the number of symbiotic algae per animal cell, however without visible bleaching. At a molecular level, we detected minimal changes in immune receptor expression patterns upon *Vibrio* exposure and most of the differentially expressed genes do not correspond to known animal genes, nor contain known conserved domains. Therefore, a completely novel immune mechanisms seems to be employed in acoels. Acoels are therefore a promising model for uncovering new immune mechanisms and for investigating the coevolution of photosymbiosis and the immune system in comparison to cnidarians.

## Methods

### Immune gene searches

Immune genes were searched in predicted transcriptomes based on the presence of relevant domains and their order (Fig. S1). The predicted proteomes were the following: *Xenoturbella bocki* (transcriptome SRX1343818), *Meara stichopi* (transcriptome SRX1343814), *Nemertoderma westbladi* (genome [179]), *Hofstenia miamia* (genome [180]), *Aphanostoma pulchra* (transcriptome SRX1343817 as *Isodiametra pulchra*), *Amphiscolops* sp. (genome [181]), *Praesagittifera naikaiensis* (genome [182]), *Symsagittifera roscoffensis* (genome [183]), and *Convolutriloba macropyga* (genome [181] and transcriptome SRX1343815). Domains were identified according to the literature (see Results), as well as through searches of established protein sequences in the NCBI Conserved Domain Database, CDD [184–186]. Alignments for the domains were downloaded from CDD and HMMER v.3.3 (www.hmmer.org [181]) was used to build Hidden Markov Model profiles (*hmmbuild*), which were used to search the predicted proteomes (*hmmsearch*). The retrieved xenacoelomorph sequences were submitted to NCBI CD Batch Search (https://www.ncbi.nlm.nih.gov/Structure/bwrpsb/bwrpsb.cgi [185]) and a sequence was kept only if it contained the domains of interests (in the correct order if relevant – see Results section). PRR sequences were considered present or absent based only on the domain structure, whereas for signalling pathway sequences we also performed phylogenetic analysis: the domain sequences identified by *hmmsearch* were aligned with MAFFT v7.526 [188,189], AMAS was used to concatenate alignments [190], and iqtree2 v2.4.0 [191] to find the best partition and model [192,193], as well as to build maximum likelihood trees, using UltraFast Bootstrap with 1000 replicates as support measure [194].

### Animals and bacteria

*Convolutriloba macropyga* Shannon & Achatz, 2007 [37] were kept in artificial sea water with salinity 36 ‰ (Classic Sea Salt by Tropic Marin, Dr. Biener GmbH, Wartenberg, Germany) at 26°C with a cycle of 10 hours darkness - 14 hours light, and they were fed twice a week with brine shrimp *Artemia*. Juveniles were collected by isolating freshly laid egg clusters. *Hofstenia miamia* Correa, 1960 [94] *we*re kept in filtered artificial sea water salinity 36 ‰ (Red Sea Salt by Red Sea Fish Pharm LTD, Eilat, Israel) at room temperature in the dark, and they were fed twice a week with brine shrimp *Artemia* sp.

*Vibrio coralliilyticus* and *Priestia megaterium* were obtained from the Jena Microbial Research Collection of the Leibniz Institute for Natural Product Research and Infection Biology, Leibniz-HKI, Jena, Germany (Strain STH00823 and STI11342, respectively). They were both grown overnight at 30±2°C with 100 rpm shaking prior to use in immune challenges. The growth media were Difco Marine Broth 2216 (Becton, Dickinson and Company, USA) for *V. coralliilyticus* and nutrient broth (DSMZ medium 1 [195]) for *P. megaterium.* A calibration curve was computed prior to immune challenge experiments as previously described [196] to correlate concentration in Colony Forming Unit (CFU)/ml to Optical Density at 600 nm (OD600).

### Immune challenges

Immune challenges were performed on *C. macropyga* adults and juveniles (0-4 days after hatching) and *H. miamia* adults. Bacteria OD600 was measured on the spectrophotometer NanoPhotometer (Implen GmbH, Munich, Germany) and bacteria concentration was calculated as 6.6·10^8^ CFUs/ml·OD600 for V. coralliily*ticus* and as 1.9·10^6^ CFUs/ml·OD600 for *P. megaterium*, according to the calibration curves. Bacteria were collected by centrifugation (1000g 5min), resuspended in filtered artificial sea water (fASW), and serially diluted. Growing medium was also centrifuged, resuspended, and diluted in the same way, as a control medium for mortality caused by media components. For immune challenges with heat-killed bacteria, the diluted bacterial suspension was incubated 5 min at 95°C with 200 rpm shaking. Animals were washed at least 3 times with fASW and isolated in 1 ml of fASW in individual wells of a 96 well plates to minimize their toxic effect [37]. The appropriate volume (25-35 µl, constant within a batch) of serially diluted bacterial suspension or growth medium was added to each well, to obtain a bacterial load of 0, 10^5^, or 10^6^ CFUs (control, low and high load, respectively). The plates were kept at animal culture conditions, but without feeding and medium exchange. They were monitored at 0, 1, 2, 4, 6, 24, and 48 hours for the 2-days assays. *C. macropyga* juveniles were monitored every day for 10 days. For long term immune challenges of *C. macropyga* adults, the animals were monitored every 2-5 days for 1 month (2 batches) or 3 months (2 batches). At each time point, animals were considered dead if not moving and showing tissue degradation, and the number of fully detached asexual progeny was counted for *C. macropyga* adults. Pictures were taken with an IPhone 14 Pro (Apple Inc., Cupertino, California, USA) mounted on a Leica M125 & (Leica Microsystems, Wetzlar, Germany).

### Dysbiosis quantification

Immune challenged *C. macropyga* adults were relaxed for 5 minutes in a 1:1 mixture of fASW and 7% MgCl_2_ and then fixed at room temperature for 30 min in 4% paraformaldehyde diluted in fASW. They were washed 5x in Phosphate Saline Buffer (PBS), incubated for 30 min in 2.5 µg/ml Hoechst 33342 in PBS, then washed twice in PBS and mounted in 70% glycerol in PBS. Whole animals were imaged with an Axiocam 503 color on an Axioscope 5 equipped with Differential Interference Contrast (DIC) (Carl Zeiss Microscopy GmbH, Jena, Germany). Chlorophyll B autofluorescence and Hoechst 33342 signal were imaged in a flat area between the mouth and the eyes with a LSM 980 confocal laser scanning microscope (Carl Zeiss Microscopy GmbH, Jena, Germany). The images were processed with a pipeline in JIPipe v5.2.0 [197]: for each channel, noise was reduced through a median filter, then brightness enhanced by dividing greyscale values by global maximum; for the algae autofluorescence channel, a difference of Gaussian was used as feature enhancer; then algae and animal nuclei were binarized automatically with a global thresholding method (Otsu for the Hoechst channel and Huang for the algae autofluorescence channel); after a morphological hole filling for the algae and distance transform watershed for both channels, the region of interests (animal nuclei and algae) were detected and counted. The length of each individual was measured on preview images for each slide with Zeiss ZEN lite microscopy software v3.11, its orientation (dorsal, ventral, lateral-dorsal) was determined based on DIC pictures.

### Gene cloning and probe synthesis

Primers were designed with Primer Blast [198] for *C. macropyga* PRR genes identified and for *V. coralliilyticus* 16S rRNA (NR_028014) (Table S6). For the latter, we selected a region that did not align to other non-Vibrio bacterial sequences retrieved with a BLASTn search [199]. Lack of aspecific amplification was checked with an *in silico* PCR tool [200] against *C. macropyga* transcriptome. Lack of aspecific hybridisation was checked *in silico* by blasting the amplicon against *C. macropyga* transcriptome within SequenceServer v2.2.0 (blastn, evalue 1e-05, sc-match 2, sc-mismatch −3, gap-open 5, gap-extend 2) [201].

Fragments of the genes were amplified by PCR from cDNA libraries obtained with SuperScript™ III First-Strand Synthesis System (Thermo Fisher Scientific, Waltham, MA, USA), inserted in pGEM-T Easy vectors (Promega, Madison, WI, USA) and transformed into competent *Escherichia coli* cells according to the manufacturer’s instructions.

Plasmids with an insert of the expected size were extracted with Plasmid Mini-Prep Kit (Jena Bioscience, Jena, Germany) and the inserts were sequenced at Eurofins Genomics Europe Shared Services GmbH (Ebersberg, Germany). Labelled antisense RNA probes were transcribed from linearized DNA using digoxigenin-11-UTP (Roche, Basel, Switzerland) with the MEGAscript T7 or SP6 kit (Thermo Fisher Scientific, Waltham, MA, USA). Probes for some scavenger receptor and C-lectin genes could not be obtained.

### Whole mount *in situ* hybridisation

Immune challenged *C. macropyga* adults were relaxed and fixed as explained above. The samples were washed 5 times in PTw (1x Phosphate Saline Buffer with 0.1% Tween-20 detergent) and stored at −20°C in methanol until further processing. Whole-mount *in situ* hybridisation was carried out as previously described [202] with the following modifications: probe concentration was 1 ng/µl and hybridisation temperature 65°C. Samples were then washed with ethanol and rehydrated in an increasing series of PTw in ethanol, before mounting them in 70% glycerol in PTw. They were imaged using an Axiocam 503 color on an Axioscope 5 with DIC (Carl Zeiss Microscopy GmbH, Jena, Germany). As a negative control, the protocol was performed without adding any probe (Fig. S7).

### Transmission electron microscopy (TEM)

Immune challenged *C. macropyga* adults were relaxed in 1:1 solution of 7% MgCl_2_ and fASW and then fixed with a mixture of 0.4% paraformaldehyde and 2.5% glutaraldehyde in 50 mM cacodylate buffer containing 23 mg/ml NaCl and 2.5 g/ml MgCl_2_. After washing in 50 mM cacodylate buffer with 23 mg/ml NaCl, the samples were post-fixed in 1% OsO_4_ in 0.1 M cacodylate buffer, washed in distilled water, and dehydrated in an increasing series of ethanol concentrations and isopropanol. They were then infiltrated in a mixture of isopropanol and Spurr resin, before embedding them in pure Spurr resin for 1 to 2 days at 60°C. *V. coralliilyticus* from liquid cultures were resuspended in fASW and subsequently processed as described above, being pelleted by centrifugation at every step (1000g, 5 min). After post fixation, they were rehydrated to bidistilled water, pelleted by centrifugation (13 000 g for 30 min), and embedded in 2% agarose, before proceeding to dehydration, infiltration, and embedding.

Ultrathin sections (70 nm) were prepared with a Leica EM UC 7 ultramicrotome (Leica Microsystems, Wetzlar, Germany) from two samples per condition (control, low-dose, high-dose) collected at 48 hours of exposure (he), one sectioned longitudinally and one transversely. In addition, transverse sections were obtained from one control sample at 1 he, two high-bacterial-load sample at 1 he, and two high-bacterial-load sample at 2 he. Ultrathin sections were mounted on slot grids and contrasted with UranyLess EM Stain (Electron Microscopy Sciences, Hatfield, PA, USA) and 3% lead citrate; the incubation was 3 min and 2 min, respectively. They were examined with Tecnai 12 transmission electron microscope (FEI Deutschland GmbH, Dreieich, Germany), equipped with a digital camera (TEMCAM FX416, TVIPS, Gauting, Germany).

### Transcriptomics and differential gene expression analysis

RNA from 2-days immune challenged *C. macropyga* adults (controls and low *V. coralliilyticus* load, 4 biological replicates of 5-10 individuals each) was extracted using TRIzol (Invitrogen, Thermo Fisher Scientific, Waltham, MA, USA) and 1-bromo-3-chloropropane (fisher scientific, Thermo Fisher Scientific, Waltham, MA, USA). The NEBNext Ultra II directional mRAN kit (New England Biolabs, Ipswich, MA, USA) was used to prepare libraries and sequencing on Illumina NovaSeq6000 platform resulted in 567.3 million total reads (average read length 151 bp). A transcriptome was assembled *de novo* with Trinity v2.15.1 [203] after quality control with fastQC (https://www.bioinformatics.babraham.ac.uk/projects/fastqc/) and trimming with fastp [204], including additional reads from *C. macropyga* juveniles processed as above (5 biological replicates, 340.4 million reads of 151 bp average length). Salmon v0.13.1 [205] was used to index the transcriptome and quantify transcript levels. Transcript level estimates, imported into R v4.3.0 [206] with the *tximport* package (design = replicate + condition, to account for paired samples) [207,208], were analysed with the *DESeq2* package [208]: principal component analysis after variance stabilizing transformation was performed with *plotPCA* and differential gene expression analysis with *DESeq*. Significantly differentially expressed genes (FDR-adjusted p-value < 0.01) were characterized by blastx search (https://blast.ncbi.nlm.nih.gov) against NCBI non-redundant protein sequences database (if default parameters did not yield any hits, compositional adjustment was set to “no adjustment” and filter low complexity regions to “unchecked”), as well as by Conserved Domain search (https://www.ncbi.nlm.nih.gov/Structure/cdd/wrpsb.cgi) [184,186,209]. If no hits or only uncharacterized proteins were retrieved, the sequences were searched against xenacoelomorph predicted proteomes used for immune gene searches with blastx v12.6.0+ [210].

### Statistical analyses and figures

Statistical analyses were carried out with R statistical software (v 4.3.0) [206]. Graphs and phylogenetic trees were produced in R v4.3.0 (Packages: *ggplot2* [211], ggtree [212] and *treeio* [213]). Figures were assembled with Adobe Illustrator. If necessary, contrast and brightness were adjusted on the whole picture with Adobe Photoshop or Fiji [214].

Survival data over time was fitted to a Cox proportional hazards mixed-effect model, with bacterial load and initial damage (accidentally caused during manipulation) as fixed explanatory variables and batch and individuals as random explanatory variables for the intercept (Table S2, *coxme* package [215]). The maximal model was simplified by sequentially eliminating all the non-significant terms and interactions, while keeping each simplified model only if equivalent to the previous more complex model (anova, α = 0.05), until a minimal adequate model was found [216,217]. The significance of fixed explanatory variables was tested with a type II Anova (*car* package [218]). When relevant, post hoc pairwise comparisons of the estimated marginal means between samples with different bacterial loads were conducted, with p value adjusted with the Bonferroni method (*emmeans* package [219]). Survival curves were plotted with ggsurvplot (*survminer* package [220]), considering fixed effects only.

For dysbiosis quantification, the ratio between the number of algae and animal nuclei was calculated for each sample from the JIPipe pipeline output. It was fitted to a linear model with bacterial load, sample length, and sample orientation (dorsal, ventral or lateral-dorsal) as interacting explanatory variables (Table S3). The model was simplified as described above. Post hoc Tukey HSD contrasts were carried out to compare bacterial loads. Animal length was fitted to a linear model (Length ∼ Bacterial.Load) and the significance of bacterial load was tested with a type II Anova, followed by post hoc Tukey HSD tests.

Asexual offspring production was fitted to a generalized linear mixed model (*glmmTMB* package [221]), with bacterial load and presence of a bud as fixed effect explanatory variables and batch as random explanatory variable for the intercept (Table S4). Data for day 1 and day 2 of exposure were analysed separately to avoid temporal pseudoreplication due to repeated measurements [216]. Different family functions were compared using Akaike’s Information Criterion (AIC) to select the more appropriate one [216,222,223]. Simplification of the maximal model was carried out as described for survival data.

## Acknowledgements

We thank all current and former members of the Hejnol working groups at the Friedrich Schiller University Jena and the University of Bergen for their support. We thank the Jena Microbial Research Collection of the Leibniz Institute for Natural Product Research and Infection Biology, Leibniz-HKI in Jena, for providing the bacteria strains used in this study. We thank the Centre for Electron Microscopy of Jena University Hospital, Friedrich Schiller University Jena, for access to their facilities for Transmission Electron Microscopy sample preparation. We thank Prof. Dr. Elisabeth Liebler-Tenorio and the Institute of Molecular Pathogenesis – Friedrich Loeffler Institute in Jena for access to the Transmission Electron Microscope and for insightful discussions. We thank Dr. Marco Groth and the Core Facility Next-Generation Sequencing of the Leibniz Institute on Aging, Fritz Lipmann Institute in Jena for their support with Illumina sequencing. We thank the University Computer Centre of the Friedrich Schiller University Jena for providing access to the HPC cluster “Draco” used to analyse transcriptomic data.

