## Supplementary material for "Loss of immune signalling pathways and increased pathogen susceptibility associated to photosymbiosis in acoels": S1 - S5

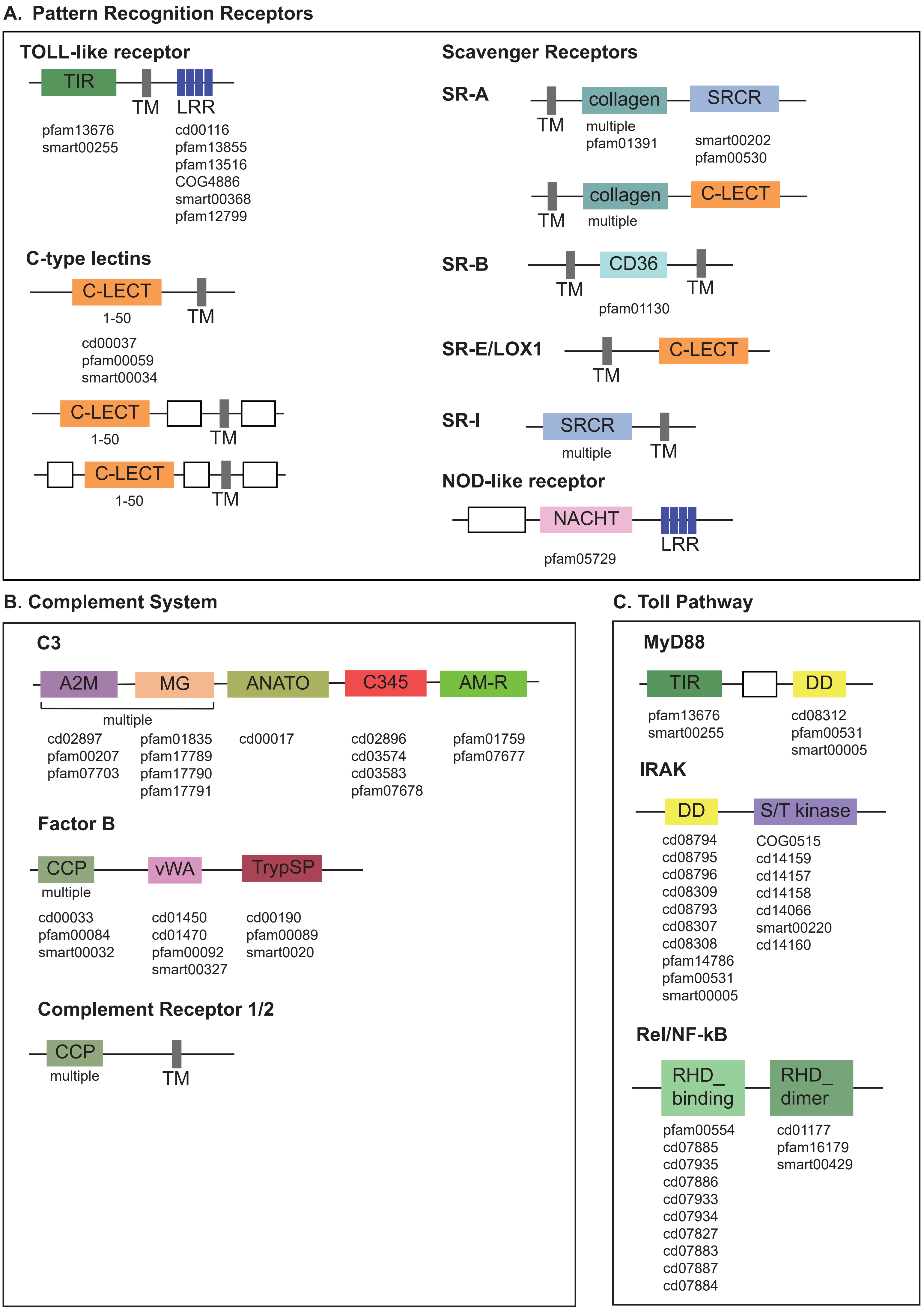
A. Pattern Recognition Receptors
TOLL-like receptor
TIR
TM
LRR
pfam13676
smart00255
cd00116
pfam13855
pfam13516
COG4886
smart00368
pfam12799
C-type lectins
C-LECT
1-50
TM
cd00037
pfam00059
smart00034
C-LECT
1-50
TM
C-LECT
1-50
TM
Scavenger Receptors
SR-A
TM
collagen
multiple
pfam01391
SRCR
smart00202
pfam00530
TM
collagen
multiple
C-LECT
SR-B
TM
CD36
pfam01130
TM
SR-E/LOX1
TM
C-LECT
SR-I
SRCR
multiple
TM
NOD-like receptor
NACHT
pfam05729
LRR
B. Complement System
C3
A2M
MG
multiple
cd02897
pfam00207
pfam07703
pfam01835
pfam17789
pfam17790
pfam17791
ANATO
cd00017
C345
cd02896
cd03574
cd03583
pfam07678
AM-R
pfam01759
pfam07677
Factor B
CCP
multiple
cd00033
pfam00084
smart00032
vWA
cd01450
cd01470
pfam00092
smart00327
TrypSP
cd00190
pfam00089
smart0020
Complement Receptor 1/2
CCP
multiple
TM
C. Toll Pathway
MyD88
TIR
pfam13676
smart00255
DD
cd08312
pfam00531
smart00005
IRAK
DD
cd08794
cd08795
cd08796
cd08309
cd08793
cd08307
cd08308
pfam14786
pfam00531
smart00005
S/T kinase
COG0515
cd14159
cd14157
cd14158
cd14066
smart00220
cd14160
Rel/NF-kB
RHD_ binding
pfam00554
cd07885
cd07935
cd07886
cd07933
cd07934
cd07827
cd07883
cd07887
cd07884
RHD_ dimer
cd01177
pfam16179
smart00429

#### Fig. S1 – Gene domain structures searched in xenacoelomorph predicted proteomes

Schematic domain structure of the genes searched in xenacoelomorph genomes and transcriptomes: **A** Pattern Recognition Receptors, **B** Toll Pathway, **C** Complement system. Under each domain name, the number of times they can be repeated (if higher than one) and the accession numbers from NCBI Conserved Domain Database used to build the hmm profiles and to filter results.

A2M = α2-macroglobulin domain; AM-R = alpha-macroglobulin binding domain; ANATO = anaphylatoxin domain; C345C = C-terminal domain specific to C3, C4, and C5; CCP = complement control protein domain; CD36 = cluster of differentiation 36; CLECT = C-type lectin domain; DD = Death Domain; IRAK = Interleukin-1 receptor-associated kinases; LRR = Leucine-rich repeat; MG = macroglobulin domain; MyD88 = Myeloid differentiation factor 88; NF-κB = Nuclear Factor-κB; RHD_bind = Rel homology DNA-binding domain; RHD_dimer = Rel homology dimerization domain (immunoglobulin-plexin-transcription domain); SR = Scavenger Receptor; SRCR = scavenger receptor cysteine rich domain; TIR = Toll/IL-1 receptor domain; TM = transmembrane region; TrypSP = trypsin-like serine protease domain; vWA = von Willebrand factor type A domain

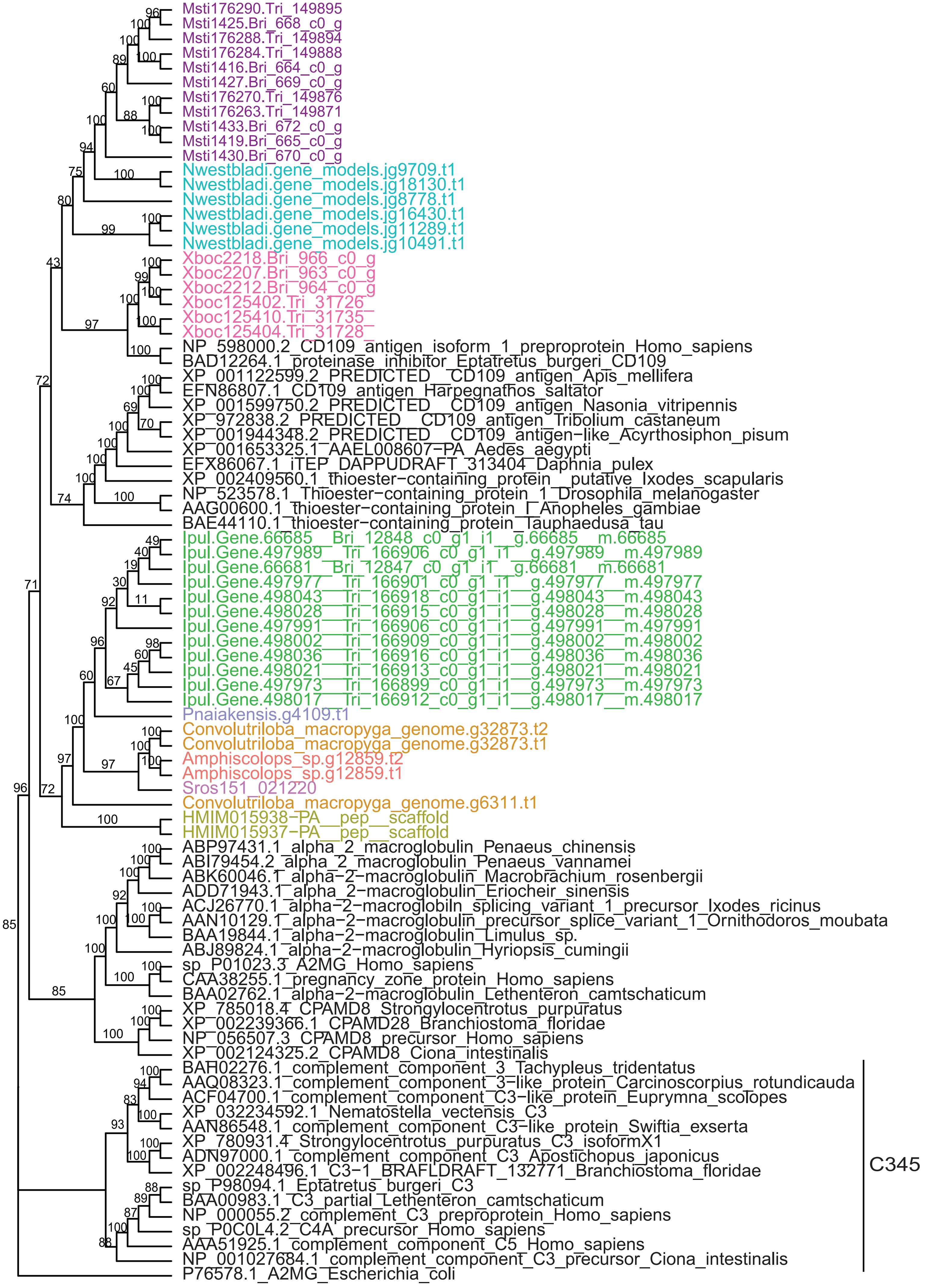

Msti176290.Tri_149895
Msti1425.Bri_668_c0_g
Msti176288.Tri_149894
Msti176284.Tri_149888
Msti1416.Bri_664_c0_g
Msti1427.Bri_669_c0_g
Msti176270.Tri_149876
Msti176263.Tri_149871
Msti1433.Bri_672_c0_g
Msti1419.Bri_665_c0_g
Msti1430.Bri_670_c0_g
Nwestbladi.gene_models.jg9709.t1
Nwestbladi.gene_models.jg18130.t1
Nwestbladi.gene_models.jg8778.t1
Nwestbladi.gene_models.jg16430.t1
Nwestbladi.gene_models.jg11289.t1
Nwestbladi.gene_models.jg10491.t1
Xboc2218.Bri_966_c0_g
Xboc2207.Bri_963_c0_g
Xboc2212.Bri_964_c0_g
Xboc125402.Tri_31726
Xboc125410.Tri_31735
Xboc125404.Tri_31728
NP_598000.2_CD109_antigen_isoform_1_preproprotein_Homo_sapiens
BAD12264.1_proteinase_inhibitor_Eptatretus_burgeri_CD109
XP_001122599.2_PREDICTED__CD109_antigen_Apis_mellifera
EFN86807.1_CD109_antigen_Harpegnathos_saltator
XP_001599750.2_PREDICTED__CD109_antigen_Nasonia_vitripennis
XP_972838.2_PREDICTED__CD109_antigen_Tribolium_castaneum
XP_001944348.2_PREDICTED__CD109_antigen−like_Acyrthosiphon_pisum
XP_001653325.1_AAEL008607−PA_Aedes_aegypti
EFX86067.1_iTEP_DAPPUDRAFT_313404_Daphnia_pulex
XP_002409560.1_thioester−containing_protein__putative_Ixodes_scapularis
NP_523578.1_Thioester−containing_protein_1_Drosophila_melanogaster
AAG00600.1_thioester−containing_protein_I_Anopheles_gambiae
BAE44110.1_thioester−containing_protein_Tauphaedusa_tau
Ipul.Gene.66685__Bri_12848_c0_g1_i1__g.66685__m.66685
Ipul.Gene.497989__Tri_166906_c0_g1_i1__g.497989__m.497989
Ipul.Gene.66681__Bri_12847_c0_g1_i1__g.66681__m.66681
Ipul.Gene.497977__Tri_166901_c0_g1_i1__g.497977__m.497977
Ipul.Gene.498043__Tri_166918_c0_g1_i1__g.498043__m.498043
Ipul.Gene.498028__Tri_166915_c0_g1_i1__g.498028__m.498028
Ipul.Gene.497991__Tri_166906_c0_g1_i1__g.497991__m.497991
Ipul.Gene.498002__Tri_166909_c0_g1_i1__g.498002__m.498002
Ipul.Gene.498036__Tri_166916_c0_g1_i1__g.498036__m.498036
Ipul.Gene.498021__Tri_166913_c0_g1_i1__g.498021__m.498021
Ipul.Gene.497973__Tri_166899_c0_g1_i1__g.497973__m.497973
Ipul.Gene.498017__Tri_166912_c0_g1_i1__g.498017__m.498017
Pnaiakensis.g4109.t1
Convolutriloba_macropyga_genome.g32873.t2
Convolutriloba_macropyga_genome.g32873.t1
Amphiscolops_sp.g12859.t2
Amphiscolops_sp.g12859.t1
Sros151_021220
Convolutriloba_macropyga_genome.g6311.t1
HMIM015938−PA__pep__scaffold
HMIM015937−PA__pep__scaffold
ABP97431.1_alpha_2_macroglobulin_Penaeus_chinensis
ABI79454.2_alpha_2_macroglobulin_Penaeus_vannamei
ABK60046.1_alpha−2−macroglobulin_Macrobrachium_rosenbergii
ADD71943.1_alpha_2−macroglobulin_Eriocheir_sinensis
ACJ26770.1_alpha−2−macroglobiln_splicing_variant_1_precursor_Ixodes_ricinus
AAN10129.1_alpha−2−macroglobulin_precursor_splice_variant_1_Ornithodoros_moubata
BAA19844.1_alpha−2−macroglobulin_Limulus_sp.
ABJ89824.1_alpha−2−macroglobulin_Hyriopsis_cumingii
sp_P01023.3_A2MG_Homo_sapiens
CAA38255.1_pregnancy_zone_protein_Homo_sapiens
BAA02762.1_alpha−2−macroglobulin_Lethenteron_camtschaticum
XP_785018.4_CPAMD8_Strongylocentrotus_purpuratus
XP_002239366.1_CPAMD28_Branchiostoma_floridae
NP_056507.3_CPAMD8_precursor_Homo_sapiens
XP_002124325.2_CPAMD8_Ciona_intestinalis
BAH02276.1_complement_component_3_Tachypleus_tridentatus
AAQ08323.1_complement_component_3−like_protein_Carcinoscorpius_rotundicauda
ACF04700.1_complement_component_C3−like_protein_Euprymna_scolopes
XP_032234592.1_Nematostella_vectensis_C3
AAN86548.1_complement_component_C3−like_protein_Swiftia_exserta
XP_780931.4_Strongylocentrotus_purpuratus_C3_isoformX1
ADN97000.1_complement_component_C3_Apostichopus_japonicus
XP_002248496.1_C3−1_BRAFLDRAFT_132771_Branchiostoma_floridae
sp_P98094.1_Eptatretus_burgeri_C3
BAA00983.1_C3_partial_Lethenteron_camtschaticum
NP_000055.2_complement_C3_preproprotein_Homo_sapiens
sp_P0C0L4.2_C4A_precursor_Homo_sapiens
AAA51925.1_complement_component_C5_Homo_sapiens
NP_001027684.1_complement_component_C3_precursor_Ciona_intestinalis
P76578.1_A2MG_Escherichia_coli
C345

#### Fig. S2 – C3 phylogeny

Maximum likelihood phylogeny of domains in C3 and related proteins. UltraFast Bootstrap support values are shown.

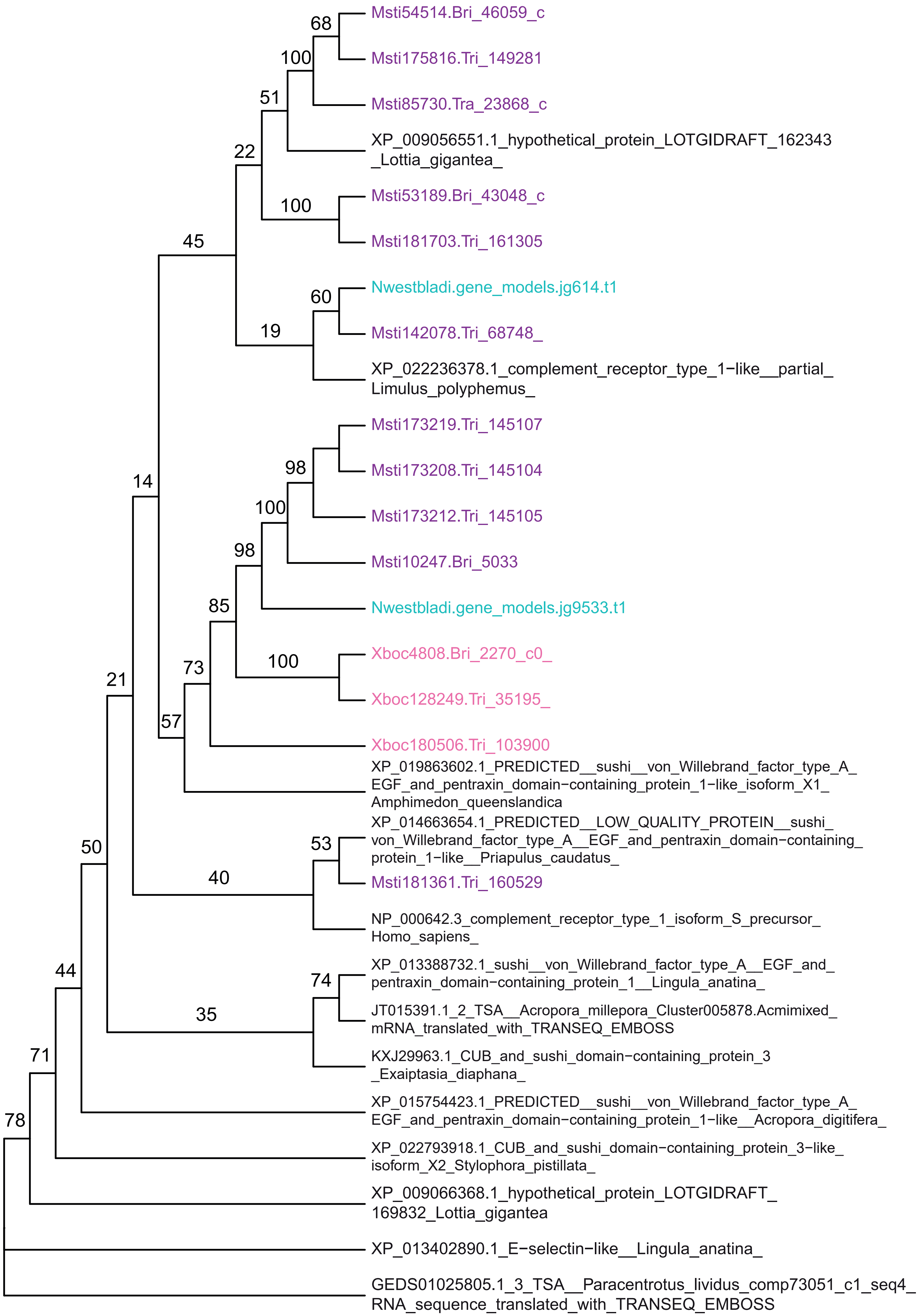
Msti54514.Bri_46059_c
Msti175816.Tri_149281
Msti85730.Tra_23868_c
XP_009056551.1_hypothetical_protein_LOTGIDRAFT_162343_Lottia_gigantea_
Msti53189.Bri_43048_c
Msti181703.Tri_161305
Nwestbladi.gene_models.jg614.t1
Msti142078.Tri_68748_
XP_022236378.1_complement_receptor_type_1−like__partial_Limulus_polyphemus_
Msti173219.Tri_145107
Msti173208.Tri_145104
Msti173212.Tri_145105
Msti10247.Bri_5033
Nwestbladi.gene_models.jg9533.t1
Xboc4808.Bri_2270_c0_
Xboc128249.Tri_35195_
Xboc180506.Tri_103900
XP_019863602.1_PREDICTED__sushi__von_Willebrand_factor_type_A_EGF_and_pentraxin_domain−containing_protein_1−like_isoform_X1_Amphimedon_queenslandica
XP_014663654.1_PREDICTED__LOW_QUALITY_PROTEIN__sushi_von_Willebrand_factor_type_A__EGF_and_pentraxin_domain−containing_protein_1−like__Priapulus_caudatus_
Msti181361.Tri_160529
NP_000642.3_complement_receptor_type_1_isoform_S_precursor_Homo_sapiens_
XP_013388732.1_sushi__von_Willebrand_factor_type_A__EGF_and_pentraxin_domain−containing_protein_1__Lingula_anatina_
JT015391.1_2_TSA__Acropora_millepora_Cluster005878.Acmimixed_mRNA_translated_with_TRANSEQ_EMBOSS
KXJ29963.1_CUB_and_sushi_domain−containing_protein_3_Exaiptasia_diaphana_
XP_015754423.1_PREDICTED__sushi__von_Willebrand_factor_type_A_EGF_and_pentraxin_domain−containing_protein_1−like__Acropora_digitifera_
XP_022793918.1_CUB_and_sushi_domain−containing_protein_3−like_isoform_X2_Stylophora_pistillata_
XP_009066368.1_hypothetical_protein_LOTGIDRAFT_169832_Lottia_gigantea
XP_013402890.1_E−selectin−like__Lingula_anatina_
GEDS01025805.1_3_TSA__Paracentrotus_lividus_comp73051_c1_seq4_RNA_sequence_translated_with_TRANSEQ_EMBOSS
68
100
51
22
100
45
60
19
98
100
14
98
85
100
73
21
57
53
50
40
44
74
35
71
78

#### Fig. S3 – CR1/2 phylogeny

Maximum likelihood phylogeny of domains in Complement receptors 1/2 and related proteins. UltraFast Bootstrap support values are shown.

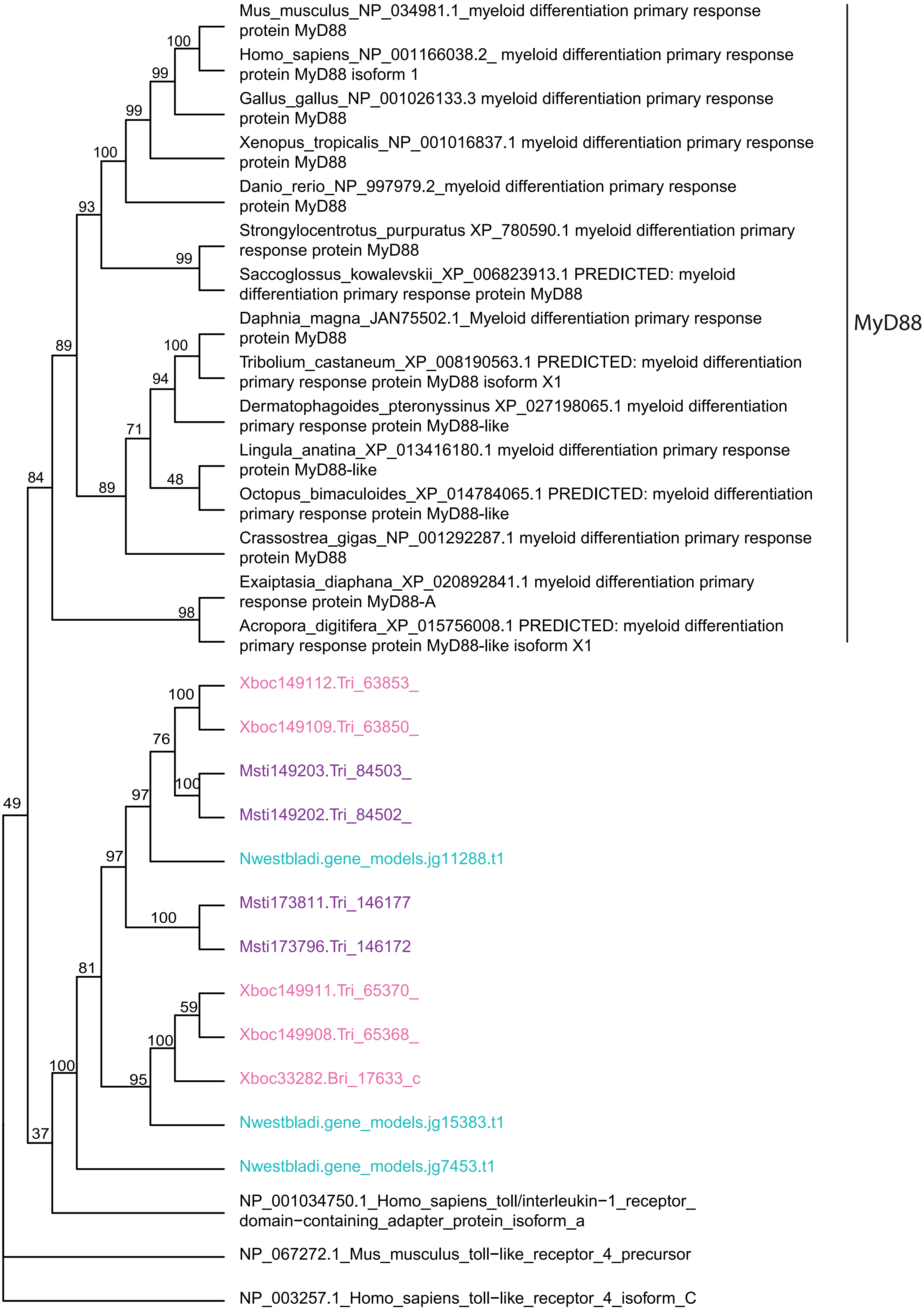
Mus_musculus_NP_034981.1_myeloid differentiation primary response protein MyD88
Homo_sapiens_NP_001166038.2_ myeloid differentiation primary response protein MyD88 isoform 1
Gallus_gallus_NP_001026133.3 myeloid differentiation primary response protein MyD88
Xenopus_tropicalis_NP_001016837.1 myeloid differentiation primary response protein MyD88
Danio_rerio_NP_997979.2_myeloid differentiation primary response protein MyD88
Strongylocentrotus_purpuratus XP_780590.1 myeloid differentiation primary response protein MyD88
Saccoglossus_kowalevskii_XP_006823913.1 PREDICTED: myeloid differentiation primary response protein MyD88
Daphnia_magna_JAN75502.1_Myeloid differentiation primary response protein MyD88
Tribolium_castaneum_XP_008190563.1 PREDICTED: myeloid differentiation primary response protein MyD88 isoform X1
Dermatophagoides_pteronyssinus XP_027198065.1 myeloid differentiation primary response protein MyD88-like
Lingula_anatina_XP_013416180.1 myeloid differentiation primary response protein MyD88-like
Octopus_bimaculoides_XP_014784065.1 PREDICTED: myeloid differentiation primary response protein MyD88-like
Crassostrea_gigas_NP_001292287.1 myeloid differentiation primary response protein MyD88
Exaiptasia_diaphana_XP_020892841.1 myeloid differentiation primary response protein MyD88-A
Acropora_digitifera_XP_015756008.1 PREDICTED: myeloid differentiation primary response protein MyD88-like isoform X1
MyD88
Xboc149112.Tri_63853_
Xboc149109.Tri_63850_
Msti149203.Tri_84503_
Msti149202.Tri_84502_
Nwestbladi.gene_models.jg11288.t1
Msti173811.Tri_146177
Msti173796.Tri_146172
Xboc149911.Tri_65370_
Xboc149908.Tri_65368_
Xboc33282.Bri_17633_c
Nwestbladi.gene_models.jg15383.t1
Nwestbladi.gene_models.jg7453.t1
NP_001034750.1_Homo_sapiens_toll/interleukin−1_receptor_domain−containing_adapter_protein_isoform_a
NP_067272.1_Mus_musculus_toll−like_receptor_4_precursor
NP_003257.1_Homo_sapiens_toll−like_receptor_4_isoform_C
100
99
99
100
93
99
89
100
94
71
48
89
84
98
100
76
100
97
97
100
81
59
100
95
100
37
49

#### Fig. – S4 MyD88 phylogeny

Maximum likelihood phylogeny of domains in MyD88 and TIR-containing proteins as outgroup. UltraFast Bootstrap support values are shown.

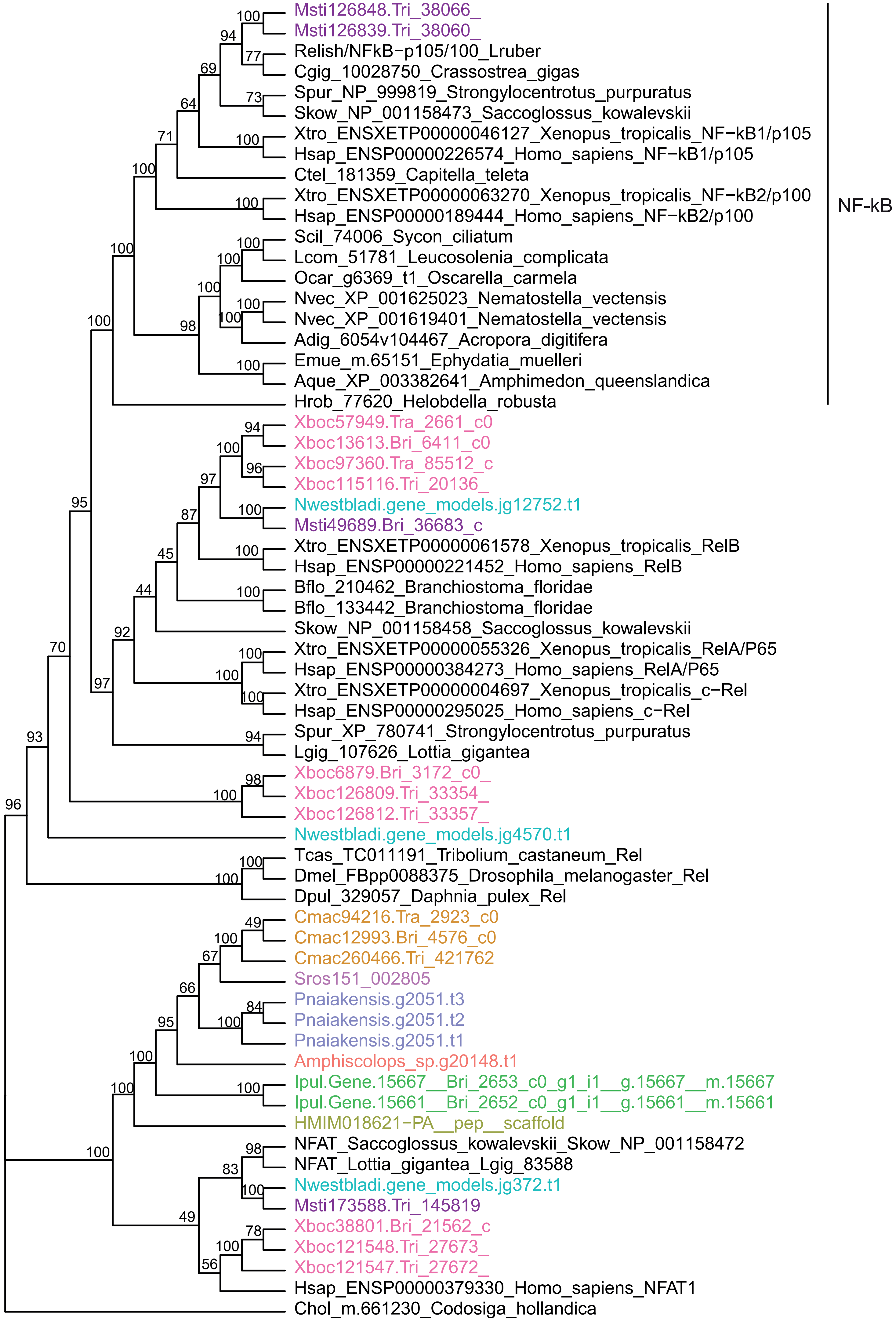
Msti126848.Tri_38066_
Msti126839.Tri_38060_
Relish/NFkB−p105/100_Lruber
Cgig_10028750_Crassostrea_gigas
Spur_NP_999819_Strongylocentrotus_purpuratus
Skow_NP_001158473_Saccoglossus_kowalevskii
Xtro_ENSXETP00000046127_Xenopus_tropicalis_NF−kB1/p105
Hsap_ENSP00000226574_Homo_sapiens_NF−kB1/p105
Ctel_181359_Capitella_teleta
Xtro_ENSXETP00000063270_Xenopus_tropicalis_NF−kB2/p100
Hsap_ENSP00000189444_Homo_sapiens_NF−kB2/p100
Scil_74006_Sycon_ciliatum
Lcom_51781_Leucosolenia_complicata
Ocar_g6369_t1_Oscarella_carmela
Nvec_XP_001625023_Nematostella_vectensis
Nvec_XP_001619401_Nematostella_vectensis
Adig_6054v104467_Acropora_digitifera
Emue_m.65151_Ephydatia_muelleri
Aque_XP_003382641_Amphimedon_queenslandica
Hrob_77620_Helobdella_robusta
NF-kB
Xboc57949.Tra_2661_c0
Xboc13613.Bri_6411_c0
Xboc97360.Tra_85512_c
Xboc115116.Tri_20136_
Nwestbladi.gene_models.jg12752.t1
Msti49689.Bri_36683_c
Xtro_ENSXETP00000061578_Xenopus_tropicalis_RelB
Hsap_ENSP00000221452_Homo_sapiens_RelB
Bflo_210462_Branchiostoma_floridae
Bflo_133442_Branchiostoma_floridae
Skow_NP_001158458_Saccoglossus_kowalevskii
Xtro_ENSXETP00000055326_Xenopus_tropicalis_RelA/P65
Hsap_ENSP00000384273_Homo_sapiens_RelA/P65
Xtro_ENSXETP00000004697_Xenopus_tropicalis_c−Rel
Hsap_ENSP00000295025_Homo_sapiens_c−Rel
Spur_XP_780741_Strongylocentrotus_purpuratus
Lgig_107626_Lottia_gigantea
Xboc6879.Bri_3172_c0_
Xboc126809.Tri_33354_
Xboc126812.Tri_33357_
Nwestbladi.gene_models.jg4570.t1
Tcas_TC011191_Tribolium_castaneum_Rel
Dmel_FBpp0088375_Drosophila_melanogaster_Rel
Dpul_329057_Daphnia_pulex_Rel
Cmac94216.Tra_2923_c0
Cmac12993.Bri_4576_c0
Cmac260466.Tri_421762
Sros151_002805
Pnaiakensis.g2051.t3
Pnaiakensis.g2051.t2
Pnaiakensis.g2051.t1
Amphiscolops_sp.g20148.t1
Ipul.Gene.15667__Bri_2653_c0_g1_i1__g.15667__m.15667
Ipul.Gene.15661__Bri_2652_c0_g1_i1__g.15661__m.15661
HMIM018621−PA__pep__scaffold
NFAT_Saccoglossus_kowalevskii_Skow_NP_001158472
NFAT_Lottia_gigantea_Lgig_83588
Nwestbladi.gene_models.jg372.t1
Msti173588.Tri_145819
Xboc38801.Bri_21562_c
Xboc121548.Tri_27673_
Xboc121547.Tri_27672_
Hsap_ENSP00000379330_Homo_sapiens_NFAT1
Chol_m.661230_Codosiga_hollandica

#### Fig. S5 – NF-κB phylogeny

Maximum likelihood phylogeny of domains in NFκB and related proteins. UltraFast Bootstrap support values are shown.

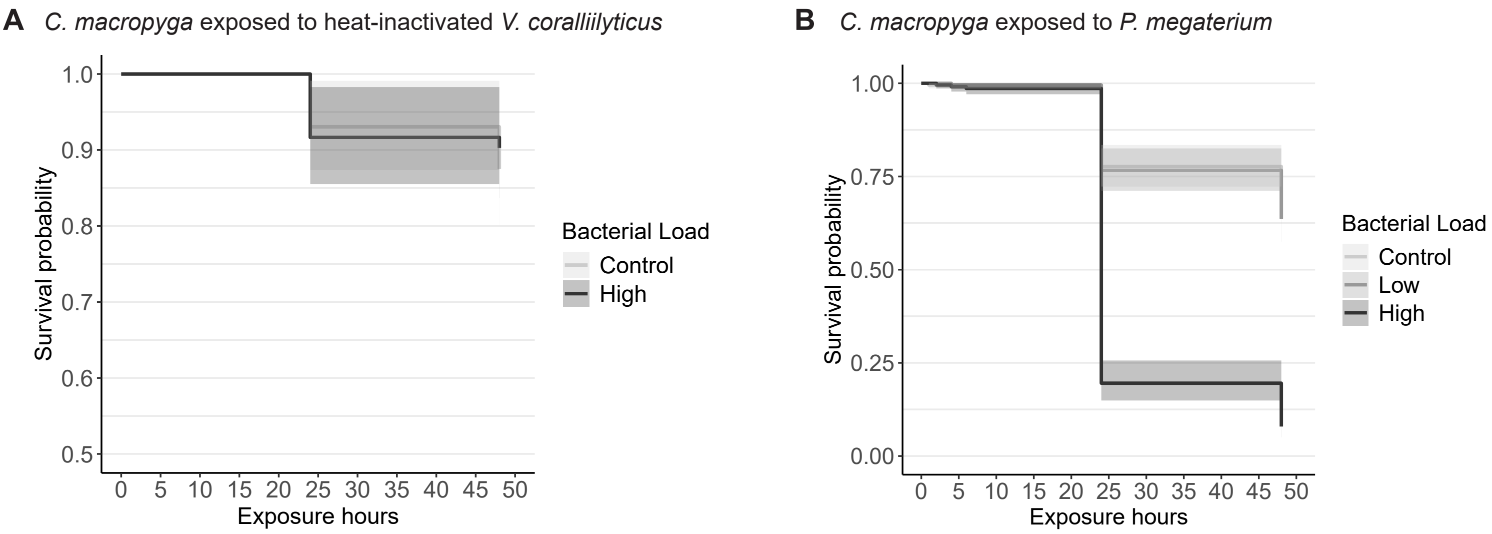


#### Fig. S6 – *C. macropyga* survival upon exposure to heat-inactivated *Vibrio coralliilyticus* and to *Priestia megaterium*

Survival curves with 95% confidence intervals for *C. macropyga* adults immune challenged with: **A** heat-inactivated *V. coralliilyticus* (minimal adequate Cox proportional hazards model Surv(last.obs,censored) ~ 1, n = 145) or **B** *P. megaterium* (minimal adequate Cox proportional hazards model Surv(last.obs,censored) ~ Bacterial.Load + (1 | batch), X2=252.3, p = 2.2e-16, n= 644).Full statistics in Table S2.

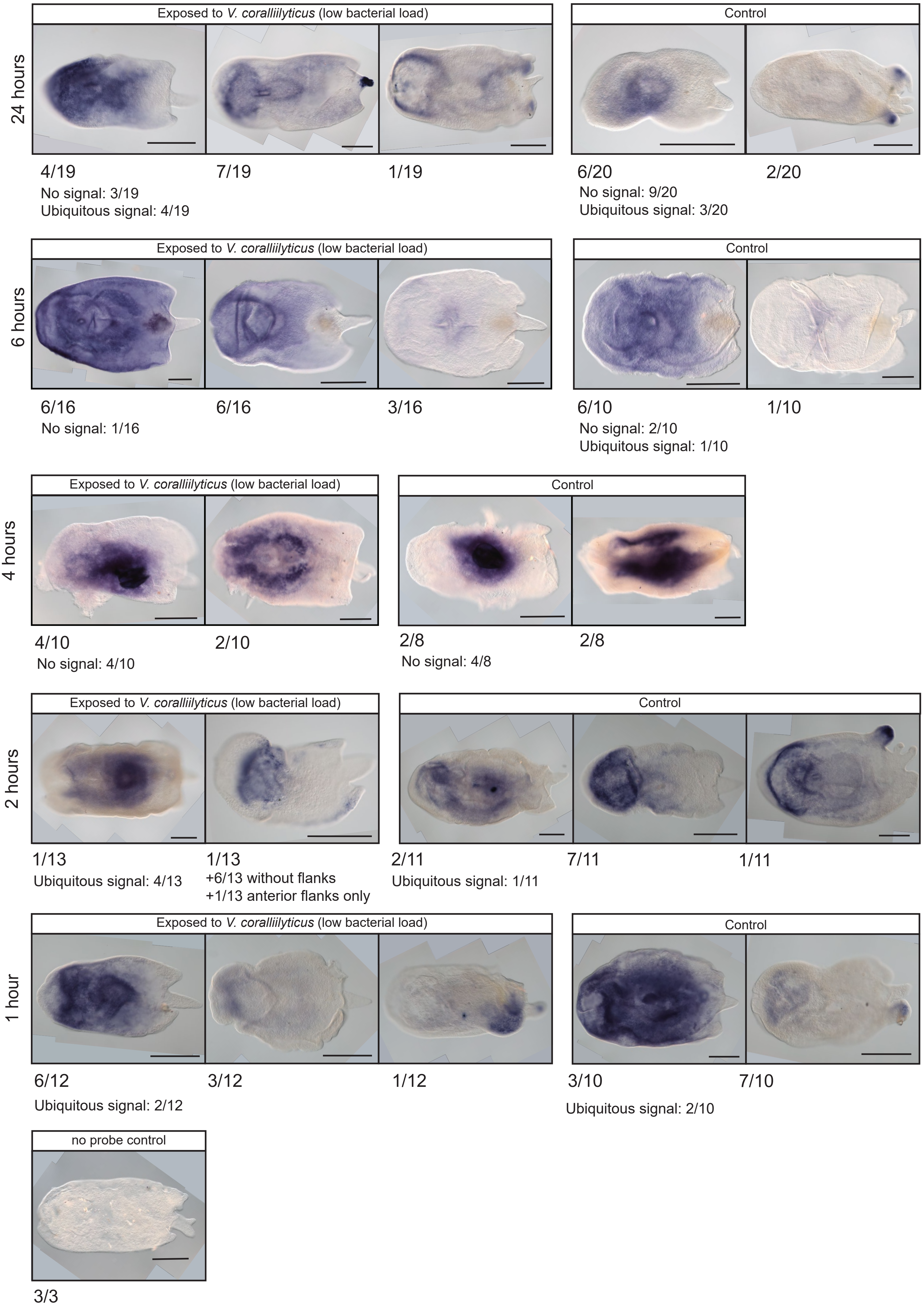
24 hours
Exposed to *V. coralliilyticus* (low bacterial load)
Control
4/19
7/19
1/19
No signal: 3/19
Ubiquitous signal: 4/19
6/20
2/20
No signal: 9/20
Ubiquitous signal: 3/20
6 hours
Exposed to *V. coralliilyticus* (low bacterial load)
Control
6/16
6/16
3/16
No signal: 1/16
6/10
1/10
No signal: 2/10
Ubiquitous signal: 1/10
4 hours
Exposed to *V. coralliilyticus* (low bacterial load)
Control
4/10
2/10
No signal: 4/10
2/8
2/8
No signal: 4/8
2 hours
Exposed to *V. coralliilyticus* (low bacterial load)
Control
1/13
Ubiquitous signal: 4/13
1/13
+6/13 without flanks
+1/13 anterior flanks only
2/11
Ubiquitous signal: 1/11
7/11
1/11
1 hour
Exposed to *V. coralliilyticus* (low bacterial load)
Control
6/12
3/12
1/12
Ubiquitous signal: 2/12
3/10
7/10
Ubiquitous signal: 2/10
no probe control
3/3

#### Fig. S7 – *In situ* hybridisation against *Vibrio coralliilyticus* 16S rRNA in *C. macropyga* at various times of bacterial exposure

RNA *in situ* hybridisation against *V. coralliilyticus* 16S in *C. macropyga* exposed to a low load of *V. coralliilyticus* or control medium for 1, 2, 4, 6, or 24 hours. Numbers indicate the ratios of individuals with the pattern shown above; dorsal view, anterior is facing left. Scale bars are 0.5 mm.

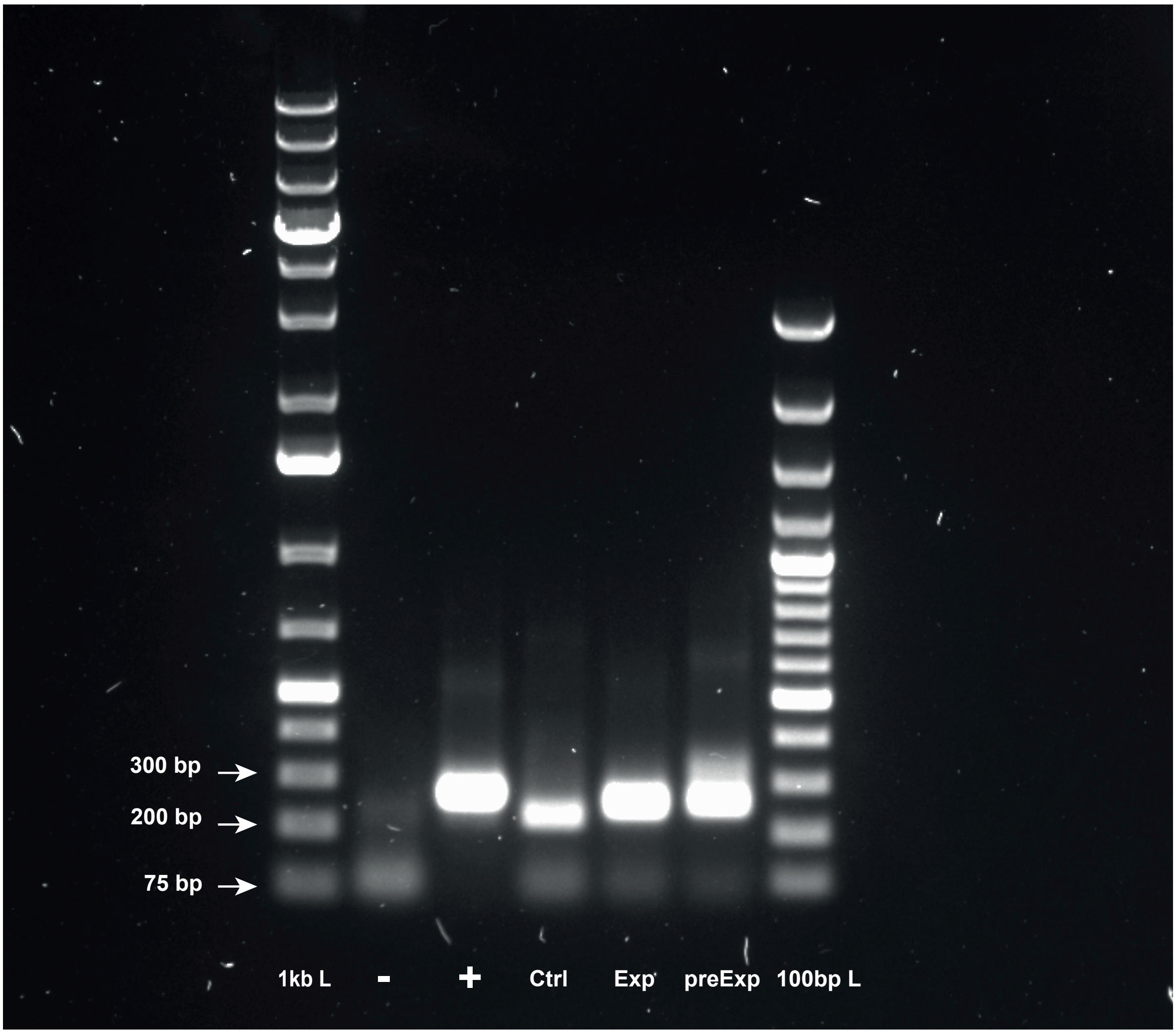


#### Fig. S8 – PCR for *V. coralliilyticus* 16S rRNA

Gel electrophoresis for PCR amplification with primers for *V. coralliilyticus* 16S from *C. macropyga* cDNA. As template: DNAse and RNAse free water as negative control (-), *V. coralliilyticus* cDNA as positive control (+), cDNA from unexposed *C. macropyga* (Ctrl), from *C. macropyga* exposed to a low dose of *V. coralliilyticus* (Exp), and *C. macropyga* cDNA synthesized before acquiring the bacterial cultures (preExp). The ladders are GeneRuler 1kb Plus DNA ladder (1kb L) and GeneRuler 100 bp Plus DNA ladder (100bp L) from Thermo Fisher Scientific (Waltham, MA, USA).

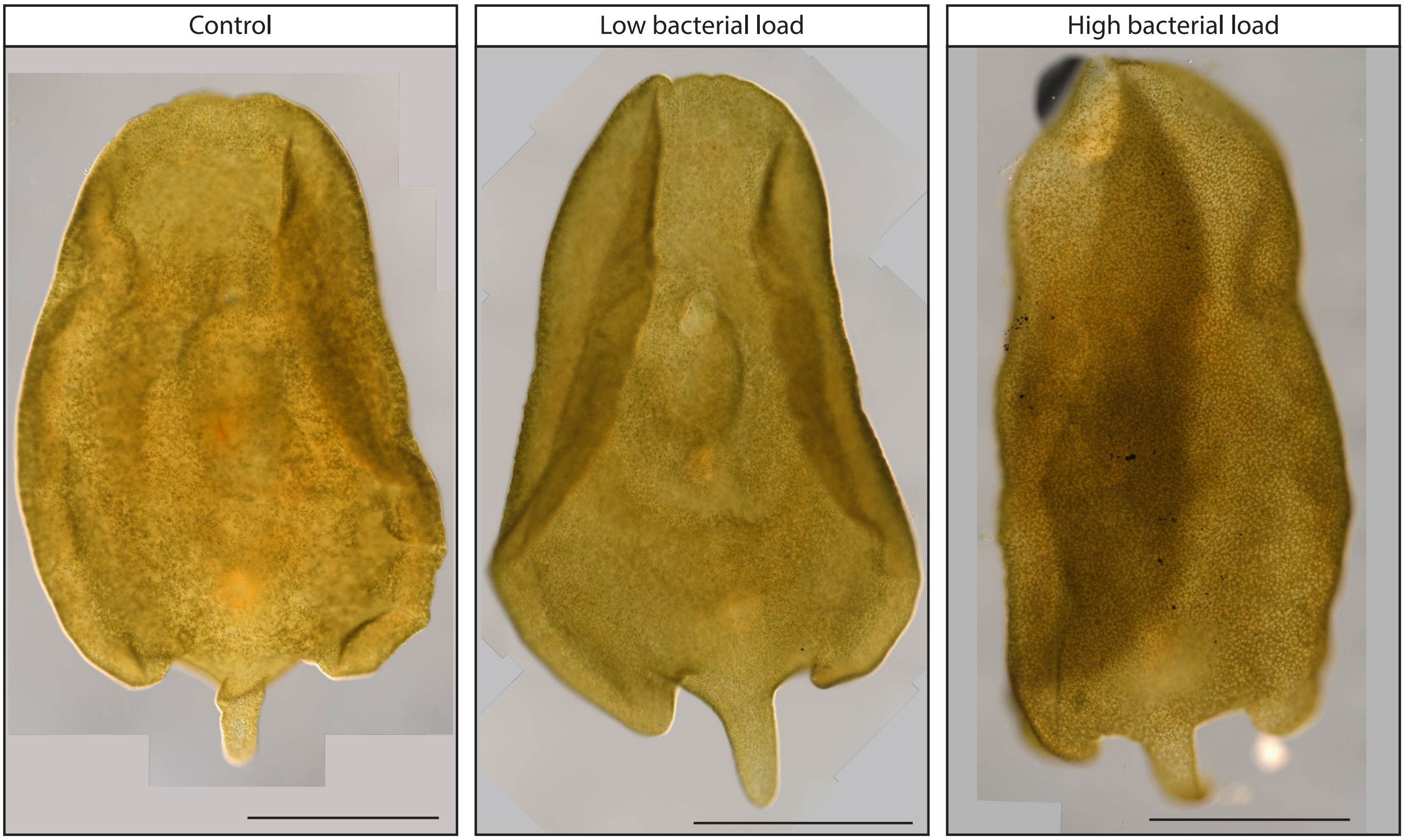


**Fig. S9 – *Convolutriloba macropyga* exposed for 14 days to *Vibrio coralliilyticus***

*C. macropyga* after 14-days immune challenges with *V. coralliilyticus* (low dose, high dose, and control), imaged with DIC after fixation and mounting. Anterior facing upwards, dorsal view.

**Video S1 – *Hofstenia miamia* expelling a sphere of tissue upon immune challenge**

### Supplementary Tables

#### Table S1

Presence and type of photosynthetic endosymbionts.

| Acoel_species | Symbionts | Reference |
|---|---|---|
| Convolutriloba_macropyga | Chlorophyta | [1] |
| Praesagittifera_naikaiensis | Chlorophyta | [2] |
| Symsagittifera_roscoffensis | Chlorophyta | [3] |
| Convolutriloba_retrogemma | Chlorophyta | [4] |
| Convolutriloba_longifissura | Chlorophyta | [5] |
| Convolutriloba_hastifera | Chlorophyta | [6] |
| Symsagittifera_corsicae | Chlorophyta | [7] |
| Convoluta_convoluta | Bacillariophyceae | [8] |
| Thalassoanaperus_tvaerminnensis | No | [9] |
| Thalassoanaperus_rubellus | No | [10] |
| Neochildia_fusca | No | [11] |
| Childia_vivipara | No | [12] |
| Childia_aculata | No | [10] |
| Childia_curinii | No | [13] |

| | | |
|---|---|---|
| Childia_crassum | No | [14] |
| Waminoa_brickneri | Dinoflagellata | [15] |
| Waminoa_litus | Dinoflagellata | [16] |
| Anaperus_sp | No | [10] |
| Hofstenia_miamia | No | [17] |
| Meara_stichopi | No | [18] |
| Nemertoderma_westbladi | No | [19] |
| Amphiscolops_sp | Dinoflagellata | [20] |
| Amphiscolops_langerhansi | Dinoflagellata | [20] |
| Haplodiscus_sp | Dinoflagellata | [21] |
| Amphiscolops_marinelliensis | Chlorophyta,Dinoflagellata | [5] |
| Amphiscolops_potocani | Chlorophyta,Dinoflagellata | [22] |
| Deuterogonaria_renei | Dinoflagellata | [23] |
| Convoluta_thela | Dinoflagellata | [23] |
| Amphiscolops_oni | Chlorophyta | [24] |
| Convoluta_cenata | Chlorophyta | [25] |
| Convoluta_sutcliffei | Dinoflagellata | [26] |
| Convoluta_enelitta | Chlorophyta | [23] |
| Symsagittifera_smaragdina | Chlorophyta | [27] |
| Symsagittifera_psammophila | Chlorophyta | [28] |
| Heterochaerus_blumi | Yes | [29] |

| | | |
|---|---|---|
| Amphiscolops_bermudensis | Dinoflagellata | [30] |
| Thalassoanaperus_biaculeatus | No | [14] |
| Thalassoanaperus_gardineri | No | [31] |
| Convoluta_boehmigi | No | [32] |
| Convoluta_cf_boehmigi | No | [32] |
| Convoluta_niphoni | Chlorophyta | [22] |
| Stomatricha_hochbergi | Chlorophyta | [33] |
| Wulguru_cuspidata | Chlorophyta | [33] |
| Daku_woorimensis | No | [33] |
| Haplogonaria_stradbrokensis | No | [33] |
| Polycanthu_torosus | No | [33] |
| Thalassoanaperus_singularis | No | [34] |
| Baltalimania_marcusi | No | [35] |
| Paratomella_rubra | No | [35] |
| Pseudaphanostoma_herringi | No | [35] |
| Aphanostoma_westbladi | No | [35] |
| Haplocelis_dichona | No | [35] |
| Aphanostoma_divae | No | [35] |
| Aphanostoma_erinae | No | [35] |
| Kuma_albiventer | No | [35] |
| Avagina_marci | No | [35] |

| | | |
|---|---|---|
| Amphiscolops_evelinae | No | [35] |
| Heterochaerus_sargassi | Dinoflagellata | [35] |
| Convoluta_henseni | Chlorophyta | [35] |
| Philactinoposthia_stylifera_brasiliensis | No | [35] |
| Kuma_asilhas | No | [35] |
| Eumecynostomum_evelinae | No | [35] |
| Aphanostoma_vexillaria | No | [35] |
| Haplogonaria_sophiae | No | [35] |
| Philactinoposthia_coneyi | No | [35] |
| Philocelis_robrochai | No | [35] |
| Amphiscolops_trifurcatus | Dinoflagellata | [36] |
| Philactinoposthia_saliens | No | [37] |
| Philactinoposthia_viridis | No | [38] |
| Philactinoposthia_brevis | No | [13] |
| Philactinoposthia_ischiae | No | [13] |
| Philactinoposthia_multipunctata | No | [13] |
| Philactinoposthia_novaecaledoniae | No | [13] |
| Aphanostoma_pulchra | No | [11] |
| Mecynostomum_tenuissimum | No | [39] |
| Childia_groenlandica | No | [39] |
| Childia_haplovata | No | [38] |

| | | |
|---|---|---|
| Notocelis_gullmarensis | No | [10] |
| Praeconvoluta_tigrina | No | [14] |
| Childia_macroposthium | No | [40] |
| Childia_submaculatum | No | [10] |
| Childia_trianguliferum | No | [10] |
| Paedomecynostomum_bruneum | No | [38] |
| Pseudmecynostomum_phoca | No | [41] |
| Mecynostomum_auritum | No | [42] |
| Paramecynostomum_diversicolor | No | [14] |
| Eumecynostomum_altitudi | No | [14] |
| Philomecynostomum_lapillum | No | [38] |
| Philomecynostomum_pictum | No | [38] |
| Aphanostoma_cuernos | No | [14] |
| Otocelis_sandara | No | [41] |
| Proaphanostoma_tenuissima | No | [43] |
| Atriofronta_polyvacuola | No | [38] |
| Pseudaphanostoma_syltensis | No | [38] |
| Actinoposthia_beklemischevi | No | [44] |
| Pseudactinoposthia_sanguineum | NA | |
| Philocelis_brueggemanni | No | [41] |
| Praeconvoluta_bruscai | No | [41] |

| | | |
|---|---|---|
| Praeconvoluta_castinea | No | [41] |
| Praeconvoluta_bocasensis | No | [14] |
| Praeconvoluta_tornuva | No | [45] |
| Praeconvoluta_collinae | No | [14] |
| Baltalimania_macrospiriferum | No | [10] |
| Aphanostoma_nicki | No | [14] |
| Aphanostoma_actuosus | No | [46] |
| Aphanostoma_virescens | No | [14] |
| Aphanostoma_bajaensis | No | [14] |
| Praeaphanostoma_schillingi | No | [14] |
| Pseudaphanostoma_smithrii | No | [41] |
| Simplicomorpha_viridis | No | [38] |
| Simplicomorpha_gigantorhabditis | No | [38] |
| Pseudohaplogonaria_rodmani | No | [47] |
| Polycanthus_torosus | No | [33] |
| Haploposthia_lactomaculata | No | [48] |
| Proporus_carolinensis | No | [34] |
| Proporus_brochii | No | [49] |
| Proporus_bermudensis | No | [50] |
| Haploposthia_rubra | No | [49] |
| Kuma_viridis | No | [51] |

| | | |
|---|---|---|
| Antrosagittifera_corallins | Dinoflagellata | [50] |
| Solenofilomorpha_crezeei | NA | |
| Endocincta_punctata | No | [52] |
| Myopea_callaeum | NA | |
| Hofsteniola_pardii | No | [53] |
| Hallangia_proporoides | No | [54] |
| Paratomella_unichaeta | No | [38] |
| Diopisthoporus_longitubus | No | [55] |
| Diopisthoporus_psammophilus | No | [10] |
| Pseudmecynostomum_bruneofilum | No | [56] |
| Oligofilomorpha_interstitiophilum | No | [56] |
| Philocelis_karlingi | No | [57] |
| Eumecynostomum_macrobursalium | No | [54] |
| Philactinoposthia_rhammifera | No | [54] |
| Baltalimania_occulta | No | [58] |
| Baltalimania_agile | No | [59] |
| Haplogonaria_minima | No | [54] |
| Haploposthia_rubropunctata | No | [49] |
| Baltalimania_fontaneti | No | [58] |
| Baltalimania_sublittoralis | No | [58] |
| Baltalimania_ylvae | No | [58] |

| | | |
|---|---|---|
| Praeconvoluta_norvegica | No | [54] |
| Xenoturbella_profunda | No | [60] |
| Xenoturbella_monstrosa | No | [60] |
| Xenoturbella_hollandorum | No | [60] |
| Xenoturbella_churro | No | [60] |
| Xenoturbella_bocki | No | [61] |
| Sterreria_lundini | No | [62] |
| Sterreria_rubra | No | [62] |
| Childia_cycloposthium | No | [57] |
| Faerlea_glomerata | No | [49] |

#### Table S2

Summary of statistical model characteristics and outputs used in the analysis of survival of *C. macropyga* adults upon immune challenge with *V. coralliilyticus* and *P. megaterium*. Model: Cox proportional hazards mixed-effects model (R *coxme* package [63]). Post-hoc comparisons: pairwise comparisons of estimated marginal means, with Bonferroni correction for P value (R *emmeans* package [64]). Degrees of freedom were infinite for all comparisons.

| | | |
|---|---|---|
| **Species, stage** | *C. macropyga* adults | *C. macropyga* adults |
| **Immune challenge agent** | *V. coralliilyticus* | *P. megaterium* |
| **Response variable** | Surv(last.obs,censored) | Surv(last.obs,censored) |
| **Number of replicates** | 4 | 3 |
| **Sample size** | 863 | 644 |
| **Maximal model** | ~ Bacterial.Load * initial.damage + (1 \| batch / id ) | ~ Bacterial.Load * initial.damage + (1 \| batch / id ) |
| **Minimal model** | ~ Bacterial.Load + (1 \| batch) | ~ Bacterial.Load + (1 \| batch) |
| **Anova (Type II)** | $X^2$=59.742, p = 1.065e-13 | $X^2$=252.3, p = 2.2e-16 |
| **Fixed effects** | | |
| **Bacterial load $10^5$ CFUs** | HR ± SE = 2.2293 ± 0.2268, z = 3.53, p = 0.000409 | HR ± SE = 0.100 ± 0.160, z = 0.00, p = 1 |
| **Bacterial load $10^6$ CFUs** | HR ± SE = 4.6465 ± 0.2104, z = 7.30, p = 2.85e-13 | HR ± SE = 6.421 ± 0.142, z = 13.13, p = 0 |
| **Random effects (Variable = intercept)** | | |
| **Batch** | SD = 0.6924, Var = 0.4794598 | SD = 0.500, Var = 0.250 |
| **Post hoc comparisons** | | |
| **Control /$10^5$ CFUs** | ratio = 0.449, SE = 0.1020, z ratio = -3.534, p = 0.0012 | ratio = 1.000, SE = 0.160, z ratio = 0.003, p = 1 |
| **Control / $10^6$ CFUs** | ratio = 0.215, SE = 0.0453, z ratio -7.301, p <.0001 | ratio = 0.156, SE = 0.0221, z ratio = -13.126, p <.0001 |
| **$10^5$ CFUs / $10^6$ CFUs** | ratio = 0.480, SE = 0.0783, z ratio = -4.500, p <.0001 | ratio = 0.156, SE = 0.0221, z ratio = -13.097, p <.0001 |

#### Table S3

Summary of statistical model characteristics and outputs used in the analysis of the ratio between algal cells and animal cells in *C. macropyga* adults upon immune challenge with *V. coralliilyticus*. Model: linear model (R *lm* function). Sample size represents the number of individuals imaged. Post hoc Tukey's tests were carried out with the R function *glht* from the *multcomp* package [65].

| | |
|---|---|
| **Species, stage** | *C. macropyga* adults |
| **Immune challenge agent** | *V. coralliilyticus* |
| **Exposure** | 48 hours |
| **Response variable** | Ratio.algae.hoechst |
| **Number of replicates** | 1 |
| **Sample size** | 74 |
| **Maximal model** | ~ Bacterial.Load * Length * Orientation |
| **Minimal model** | ~ Bacterial.Load |
| **Anova (Type II)** | F = 5.5069, p = 0.006961 |
| **Tukey HSD Contrasts** | |
| **Low bact. load - control** | t = -1.348, p = 0.37601 |
| **High bact. load - control** | t = -3.298, p = 0.00519 |
| **High bact. load – low bact. load** | t = -1.861, p = 0.16080 |

#### Table S4

Summary of statistical model characteristics and outputs used in the analysis of asexual progeny release of *C. macropyga* adults upon immune challenge with *V. coralliilyticus*. Model: generalized linear mixed-effects model (R *glmTMB* package [66]). Estimated marginal means (EMMs) obtained with R *emmeans* package (specs = ~ bud, type = "response") [64]. Sample size corresponds to the number of alive individuals at the corresponding day.

| | | |
|---|---|---|
| **Species, stage** | *C. macropyga* adults | *C. macropyga* adults |
| **Immune challenge agent** | *V. coralliilyticus* | *V. coralliilyticus* |
| **Exposure day** | Day 1 | Day 2 |
| **Response variable** | progeny.released | progeny.released |
| **Distribution function** | Poisson | Poisson |
| **Number of replicates** | 4 | 4 |
| **Sample size** | 813 | 669 |
| **Maximal model** | ~ Bacterial.Load * bud + ( 1 \| batch) | ~ Bacterial.Load * bud + ( 1 \| batch) |
| **Minimal model** | ~ bud + (1 \| batch) | ~ bud |
| **Anova (Type II)** | $X^2$ = 24.684, p = 6.752e-07 | $X^2$ = 18.512, p = 1.688e-05 |
| **Fixed effects (EMMs)** | | |
| **Bud absent** | rate ± SD = 0.171 ± 0.0266 | rate ± SD = 0.108 ± 0.0136 |
| **Bud present** | rate ± SD = 0.422 ± 0.0851 | rate ± SD = 0.304 ± 0.0620 |
| **Random effects (Variable = intercept)** | | |
| **Batch** | SD = 0.252, Var = 0.06351 | - |

#### Table S5

Differentially expressed genes between *C. macropyga* adults immune challenged a low *V. coralliilyticus* load and controls at 2 days of exposure.

| baseMean | log2FoldChange | lfcSE | pvalue | padj | BlastTopHits | ConservedDomains | CD_AN | Hits_xenacoelomorph_db |
|---|---|---|---|---|---|---|---|---|
| 682.0843165 | -1.62234 | 0.243572404 | 4.35E-13 | 1.52E-08 | Bacterial protein (Chlamydiales, Mycoplasma, Clavispora, Argiope, Trichomonas - not Vibrio) | Mplasa_alph_rch super family - CwlO1 super family | TIGR04523 - COG38833 | |
| 361.1305306 | -1.53312 | 0.271027905 | 6.57E-10 | 1.15E-05 | secreted RxLR effector protein 161-like Xenia sp. Carnegie-2017 | RNase_H_like super family | cd09272 | |
| 6169.828611 | 0.789815 | 0.142559871 | 1.24E-09 | 1.44E-05 | Uncharacterized S. roscoffensis protein | NA | | Cmac, Pnai, Sros |
| 473.5593284 | -1.89E-05 | 0.001442744 | 2.77E-09 | 1.93E-05 | NA | NA | | Cmac |
| 409.4232391 | -1.5363 | 0.28337625 | 3.04E-09 | 1.93E-05 | NA | NA | | NA |

| | | | | | | | | |
|---|---|---|---|---|---|---|---|---|
| 443.1927491 | 0.965198 | 0.179831485 | 3.31E-09 | 1.93E-05 | Serine dehydratase | Trp-synth-beta_II super family | cl00342 | |
| 5747.800264 | 0.716368 | 0.134608318 | 4.19E-09 | 2.10E-05 | Uncharacterized S. roscoffensis protein | NA | | Cmac |
| 3196.515604 | -0.71383 | 0.136093117 | 5.67E-09 | 2.48E-05 | Uncharacterized protein from Ancylostoma, Phytomonas, Trypanosoma, ice nucleation protein from Anopheles, extensis-like from Saccoglossus, tartrate-sensitive acid phosphatase from Leishmania | NA | | Cmac, Sros |
| 299.6581054 | -1.69692 | 0.325026974 | 7.05E-09 | 2.74E-05 | Serine protease | Secreted trypsin-like serine protease - Tryp_SPc | COG5640 - cl21584 - cl44276 | |

| | | | | | | | | |
|---|---|---|---|---|---|---|---|---|
| | | | | | | super family - Secreted trypsin-like serine protease | | |
| 870.9337797 | -0.9065 | 0.177274177 | 1.30E-08 | 4.55E-05 | NA | NA | | Cmac |
| 328.9283974 | -1.57981 | 0.319044988 | 2.65E-08 | 8.44E-05 | C-type lectin | Lectin_c | pfam00059 | |
| 30856.58096 | -0.7624 | 0.154224011 | 3.08E-08 | 8.99E-05 | dynein regulatory complex subunit 4-like | NA | | |
| 1572.103548 | -0.7714 | 0.164670727 | 1.03E-07 | 0.000277 | NA | NA | | Cmac |
| 140.9937225 | -1.83371 | 0.401065818 | 1.75E-07 | 0.000437 | Unnamed protein from Pseudomonas | NA | | NA |
| 413.9812218 | -1.58E-05 | 0.001442726 | 3.90E-07 | 0.000911 | Serine protease | Tryp_SPc | cl21584 | |
| 437.1652343 | -1.35752 | 0.314519575 | 5.15E-07 | 0.001126 | NA | NA | | Cmac |
| 310.1880361 | 0.827524 | 0.194152735 | 8.01E-07 | 0.001649 | WXG100 family type VII secretion target | NA | | |
| 51.53777295 | -2.10392 | 0.503391713 | 9.91E-07 | 0.001828 | NA | NA | | NA |
| 439.9775298 | -0.84157 | 0.201458325 | 9.93E-07 | 0.001828 | Unnamed protein from Closterium sp (single hit) | NA | | |
| 388.2579093 | -0.98282 | 0.240630072 | 1.48E-06 | 0.002587 | NA | NA | | Cmac |

| | | | | | | | | |
|---|---|---|---|---|---|---|---|---|
| 6265.722089 | 0.527218 | 0.12912357 | 1.63E-06 | 0.002717 | Uncharacterized proteins as best hits, GTP-binding A, DHH, sonic hedgehog | Hint module - MAC/Perforin domain - predicted GTPase | pfam01079 - pfam0183 - COG3596 | |
| 263.7528007 | 1.00564 | 0.251375436 | 1.99E-06 | 0.003158 | Uncharacterized proteins from S. roscoffnesis and cnidarians | NA | | Amph, Cmac, Hmia, Apul, Pnai, Sros |
| 2099.699433 | -0.62131 | 0.155747389 | 2.50E-06 | 0.003809 | Mucin-2-like, adhesive plaque matrix protein, trophinin-like, FP1V1 protein | Nidogen-like - Topoisomerase II-associated protein PAT1 | cl02648 - cl37801 | |
| 129.0383752 | -1.46384 | 0.370058274 | 2.91E-06 | 0.004085 | NA | NA | | Cmac |
| 1605.481256 | -0.67144 | 0.172212013 | 2.92E-06 | 0.004085 | Carbonic anhydrase | Eukaryotic-type carbonic anhydrase | pfam00194 - cl47513 | |
| 196.345473 | -1.33397 | 0.354103903 | 5.10E-06 | 0.00687 | Serine protease | Secreted trypsin-like serine protease | COG5640 | |

| | | | | | | | | |
|---|---|---|---|---|---|---|---|---|
| 308.959335 | -1.244 | 0.329284597 | 5.57E-06 | 0.007221 | NA | NA | | Cmac |
| 9573.467933 | 0.59994 | 0.165964647 | 7.96E-06 | 0.009607 | Uncharacterized S. roscoffensis protein | NA | | Cmac, Pnai, Sros |
| 3864.037974 | 0.589319 | 0.160379047 | 7.88E-06 | 0.009607 | Transient receptor potential cation channel subfamily M | TRPV - LSDAT_euk | TIGR00870 - pfam18139 | |

#### Table S6

Primers used for cloning.

| gene | ID | for_primer | rev_primer | in_silico_amplicon |
|---|---|---|---|---|
| Vcor16S | NR_028014 | TAACACATGCAAGTCGAGCGG | AGACCAGCTAGGGATCGTCG | TAACACATGCAAGTCGAGCGGAAACGAGTTGTCTGAACCTTCGGGGAACGATAACGGCGTCGAGCGGCGGACGGGTGAGTAATGCCTGGGAAATTGCCCTGATGTGGGGGATAACCATTGGAAACGATGGCTAATACCGCATAATAGCTTCGGCTCAAAGAGGGGGACCTTCGGGCCTCTCGCGTCAGGATATGCCCAGGTGGGATTAGCTAGTTGGTGAGGTAATGGCTCACCAAGGCGACGATCCCTAGCTGGTCT |
| C-lectin 1 | Cmac.Gene.115445::Tra_24119_c0_g1_i1::g.115445::m.115445 | CGAGAAGTATCTCCGACCCC | TGGTCACAAAAATTCACCGCC | CGAGAAGTATCTCCGACCCCTGGAACATCTAATGATGCTATTGAGGTCAAAGGACAAGACAACATTGTGACTGAGCCGCGGAGAACGATCAACTCTCCTGTTTATTATATCGATAATCTAGTGCGATCTGGCAGTTCTAGACCTCATTTCGAGGACAGTACCCAATATTCTGCTTCGGCTGCAGCATCAGATGCAAATTTACTCAAAAACTATCAATATGATTTTCCAAACTCGTGCATTTTGTACACTTTGTTAGCGATGATCGGTGTGACTGTCT |

| | | | | |
|---|---|---|---|---|
| | | | | TCGCAGTAGCATTTAATTTAGCTTGTCCTATTTTTGTGTTGAACCGACTGGACGAGTTAGTATCTGAAGAGGTTGCTGATGTTTTAAAAGAGCAAGATCTCGACACGCTCAATTGTGCTGCCGATTGGCTGCCATATCGTCAGAATTGCTATCGGAAATACAGAAGCAAGACAGCAAGAAATTATTCTAGGGCGGTGAATTTTTGTGACCA |
| C-lectin 2 | Cmac.Gene.63052::Bri_40559_c0_g1_i1::g.63052::m.63052 | CATTGAGAAGGCCAAATATGCG | TTTCTCGTGTCGCCTGTCG | CATTGAGAAGGCCAAATATGCGATAGTATCAGCTTCATGTGAGCAGCATGGTGGTGTTATATATGGTCGAAAGTGCTTTTCAATTTTGTGGAACAAAGCGTACAACAGAACTTTTCAAGCAGCCAAACAGTTTTGTAAGTCTCAGAACCAAACCCTCGCCTACCCATCAAACAAAGGTGAACTTCACTTTATGACCAAGTTCATGTGGAGTTTTGAGCACTTAACGCCGCATGTTGGGTTCGAGCGGTCGGGCGATAGTTCTGACTTCGTTTCAGTCGACGGGCGAGTTAAACTGAATCCGACGTCTAATATGTGGGATGTAGGGTTCCCGGCTGAGGGACGAGGTCGGTTTGGATTCCTGAGGATGGACACGACAGGCGACACGAGAAA |
| C-lectin 3 | Cmac.Gene.272337::Tri_454578_c0_g1_i1::g.272337::m.272337 | ACTCTCATCTCGCGTTTCCC | AGGAGTGGAACAGTCTTGCG | ACTCTCATCTCGCGTTTCCCAGAGACGCTCTAGAGATGGCTGTCCTCAAATCAATGGCTCCTTTTGATCCTGCTCAAAATAACCCGGTAACGTTTCTTGGCCTTAGCCGCACAAGCGAAGCCTTGAATGACTGGCAATTTCACGACAGCACAACCTACAACATTTCTCAGAAACTAGACGAGCTTTGGGACATTGGGGAGCCAAACCCAGCTGGCAACTGTGCTGCATTCATAGTCGGTTCCGCCCATCACAACAAAGTGGTCACGCAAGACTGTTCCACTCCT |
| NLR | Cmac.Gene.27073::Bri_11364_c0_g1_i1:: | CGGTCCTGCTGGTATTGGG | AAGCCAGTCCAATTTGTCAGC | CGGTCCTGCTGGTATTGGGAAGTCGACGTTATTGGGACAAATTGCGTGCAGGTTTTTGAAAGGAGATTTGCCTAAGTTCCAGAATGTTATTTTGCTCACTTGTCGAAAATTGAACCAATTCGGAGGGAAGAAAATGACTTTTGAGCATTTCATTCAAAGCTATGGTTACACTGTCGACTTTGACTTCAGCTCGCCGCATTTTGCGGATCAGTTGTTTTTGTT |

| | | | | |
|---|---|---|---|---|
| | g.27073::m.27073 | | | CGATGGTTTCGAGGAGTACAATCGAAGGTGGACATTCGGAAAGGAGATGACTGGAGTTGAGGGCCGTGTTTCCGTTGATTCCTCAGAAATGACAACTGATGAATGGATCCGCTACTTAATGTGCTCAAATGGTAAAAGTGTTGTGATTGCAGCGCGACCAGGGTCCCTTAGATTCTTGAAGGATTACCCTTACAAACTGAGACTCAGGGTTCTTGGTTTCAGTGATCAAATGATCCGAGAATTCTTCAAAATGCATGGCAGTGCTGACAAATTGGACTGGCTT |
| SR-B 1 | Cmac.Gene.8399::Bri_3021_c0_g1_i1::g.8399::m.8399 | CGTCCTTAGCTGATGGCTCG | ACACTCGTCCTTTTCTCCGC | CGTCCTTAGCTGATGGCTCGATGGTCTATGACGAATGGCAGGCACCCGACTACCCAATCTATATGAAGTATTACATGTACACATACACCAATGCACAAGACTTCGTCGATGGCACAGATCCTAAGCCAGAGGTAGAACAACTCGGCGTCCTATCTTATCGAGAGACACAAGTGAAGTTTGATCTTGAGTACGTGGATGACGACAAAAGAGTGATCTACTACAATAACAAATCGTATGTCTTCGATGAAGAAACATCAACAATGGCCGAAAATGACACAATTGTAGTTCCTAACATTCTTCTCTTTACGATTCTGGCGACATATCAAGAGACGCTTAATTGGGGCTTCATCAAGTTTTTTGCTAACCGTGACTTCAATGGAGGTGATCCTGACTACGTAAACGCAGTGTCTCCATTCTACATGGTGGAAGCGGGCGACTTTCTCTGGGGATATAACGATAATCTCTTTCTAGAGCAATTGCATGGATTGGATGCCGAAGTGCCTACAAAATTTGGACTCCAGGTGAACAATACAAATGATGGTCAATACGAGATATACACAGGGAAAGGCGATGACAACGACAAGCGAGGAATTGTTGAGAAATGGGACAATTTGACTGCTCTTACTTTCTGGTACAGTGACACGTGCAATATGATCAATGGCACAGATGGCACTATATTCCCTGTGGATGTCCCAAAAGATGAAAGACTGTATGTGTTCAATACTCAAATTTGCAGAAGTATCTTCTTAGAAGCTGAGGAAGATCAAGAGATAGAAGGAATTGATACTACTAAGTTTGTGGCTCCTAGTGAAGTTTTCGAAGTGGACTTTGAAGATAATGTTGGCTATTGTACGGGCGGAGAAAAGGACGAGTGT |

| | | | | |
|---|---|---|---|---|
| SR-B 2 | Cmac.Gene.244825::Tri_389305_c0_g1_i1::g.244825::m.244825 | TGGACTCACAATCCTGTTCGG | AAAACCTCTCCGTCAGTGCC | TGGACTCACAATCCTGTTCGGCTACCCTTCATTCTACAATAAACAAACATCCCTAGTAAGAGGTAGCCAGACGTACCAAGAGTGGGTGAAGCCAGATCTGCCTGTGTACATGCGGTACCACATGTTCAATTACACAAATGCCGACAAGTTCATTCAGAATGTAGACGAAAAGCCGATTTTGGAAGAAATTGGCATTCTGTCATACAGAGAGTACGAAGAGAAACACGATGTAGAATACGTGAAGTCTGGTGACATTGTGAGATATGTGAGCAACAGGACATATGTATTCGACCCTTTCACTTCTACCATGAATGAGTCTGACATCGTTACAGTCCCAAACATTGTCATGTTTACTGTGTTGGCATCAGCGCCTGCTCTGGAGAGAGGAGTTGTGGAAGTAAAATTTGACGAGTGGGCTCTGGAATTCAATCATCCAGACCTGCAGAATGCAGTCTCCCCGTTTTATCACGTGGAAGCGGGTAACTTCTTGTGGGGTTATGAAAACAACTCGCTCCTTGTCAAGTTGGAAGAGTTGATAGGAGGACCGACTAGTTTCGGATTACAGTCTAACAACTCAAATGACGGGCCTTACGCCGTGCTAACCGGGAAGGACAATTTGAACGAGAAAAGAGGCAAGATTATCCAGTGGTACAACATGCAGTCATTGCCTTTCTACAGTAGCAAGTACGCTAACATGATCAACGGCACTGACGGAGAGGTTTT |
